# cMagnetic Resonance Reveals Radical Stability in Native Biofilms and its Role in Bacterial Resistance

**DOI:** 10.1101/2025.07.21.665160

**Authors:** Alysia Mandato, Hannah Hunter, Sadhana Srinivasa, Chang-Hyeock Byeon, Abdulkadir Tunc, Meghan K. Wells, Faith J. Scott, Frederic Mentink-Vigier, Wook Kim, Sunil Saxena, Ümit Akbey

**Affiliations:** Department of Chemistry, University of Pittsburgh, Pittsburgh, PA 15213, USA; Department of Biological Sciences, Duquesne University, Pittsburgh, PA 15282, US; Department of Structural Biology, School of Medicine, University of Pittsburgh, Pittsburgh, PA 15261, USA; National High Magnetic Field Laboratory Florida State University, allahassee, FL 32310, USA

**Keywords:** bacterial biofilm, EPR spectroscopy, nitroxide, NMR spectroscopy, *Pseudomonas fluorescens*

## Abstract

Bacterial biofilms exhibit enhanced antimicrobial resistance, yet the mechanisms determining molecular transport and reactivity within these complex communities remain poorly understood. Here, we combine electron paramagnetic resonance (EPR), solid-state NMR (ssNMR), and dynamic nuclear polarization (DNP) ssNMR to determine how native *Pseudomonas fluorescens* Pf0-1 colony biofilms regulate nitroxide radicals. EPR measurements reveal that radical reduction depends on biofilm morphology, hydration, and composition of extracellular matrix (ECM). Wild-type biofilms exhibit slower nitroxide reduction than planktonic cells, while isolated ECM and dehydrated biofilms show no radical reduction, indicating that bacterial cells primarily drive radical reduction. Complementary ssNMR measurements identify polysaccharides and lipids within the ECM as primary interaction sites for nitroxide radicals. DNP-enhanced ssNMR further reveals compositional differences across biofilm morphologies, with increasingly polysaccharide-rich ECM environments correlating with slower reduction kinetics. These findings support a mechanism in which the ECM acts as a diffusion barrier and selective interaction region for nitroxide radicals to regulate cellular penetration. This work showcases an integrated magnetic resonance approach that provides molecular insight into how biofilm structure determines the fate of toxic redox-active small molecules. We also set the stage for high-sensitivity measurements of structure-function relationships in these medically relevant assemblies.

## Introduction

Many species across the bacterial taxa naturally form biofilms that protect the community members from diverse environmental stressors. During biofilm formation, individual cells first attach onto surfaces or auto-aggregate then surround themselves in a self-produced extracellular matrix (ECM) that is composed of secreted polysaccharides, extracellular DNA, lipids, proteins, and other molecules.^[1, 2]^ Within this matrix, cells coexist in a coordinated community that enables them to share nutrients, exchange signals, and resist environmental stress. Compared to free-floating planktonic bacteria, biofilm-associated cells have enhanced resistance to antimicrobial agents, partly because the ECM acts as a protective and diffusion-limiting barrier. Consequently, biofilm cells are up to 1,000 times more tolerant to antimicrobial drugs than planktonic cells.^[1-3]^ Such biofilm-protected bacteria, including *Staphylococcus aureus* and *Pseudomonas aeruginosa*, cause approximately 80% of all chronic infections, particularly in cystic fibrosis cases, and are among the most critical human pathogens recognized by the World Health Organization.^[4-8]^ Current strategies to counter biofilm-mediated antimicrobial resistance focus on surface treatments that disrupt bacterial adhesion, degrade the secreted polysaccharides, or block quorum sensing to prevent biofilm establishment. However, treatment efficacy remains low, as biofilms display high levels of diversity between species, individual samples, and can even change over time as the matrix evolves in response to stressors and antibiotics.^[9, 10]^ Despite growing interest in biofilm-targeting treatments, little is known about how native biofilm environments regulate the stability, reduction, and diffusion of small molecules within the ECM. To date, there has been no systematic study on the relationship between ECM structural composition and its impact on antimicrobial resistance.

Importantly, oxidative and nitrosative stressors are known to disrupt bacteria and biofilm formation.^[11, 12]^ In particular, nitroxide radicals (acting as nitric oxide mimics) have emerged as promising antibiofilm agents, capable of disrupting biofilm formation and promoting dispersal.^[13]^ For example, low concentrations of nitric oxide induce dispersal of the biofilm into free-floating planktonic bacteria that are significantly more susceptible to antimicrobials than the biofilm.^[11, 14-17]^ Nitroxide compounds and nitroxide-conjugated antibiotics have therefore been developed as a potential method for fighting biofilm-associated infections.^[18-22]^ Moreover, elevated levels of intracellular nitric oxide induce bacterial toxicity.^[11, 23, 24]^ Nitroxides have been shown to be effective against both Gram-negative and Gram-positive biofilm-producing bacteria while simultaneously being non-toxic to normal human cells.^[25, 26]^ Given that nitroxides function through redox-active radical states, their stability and therapeutic behavior likely depend strongly on the local environment within the biofilm and ECM. These mechanisms of radical reduction and stability remain poorly understood.

To investigate the relationship between nitroxide molecules and the biofilm, herein we focus on *Pseudomonas fluorescens* (Pf0-1). This organism displays remarkable multicellular behaviors and rapidly adapts its ECM composition in response to environmental stress. Previous works have demonstrated that wild-type (WT) Pf0-1 can diversify into distinct morphotypes, including mucoid and dry variants, which cooperate and spatially self-organize to form complex collective structures with a division-of-labor-like behavior.^[27]^ These phenotypic variants exhibit striking differences in ECM production, colony morphology, spreading dynamics, and mechanical stability. Therefore, Pf0-1 is an excellent system for investigating how differences in chemical composition and organization govern the behavior of small molecules within native biofilms.

Despite these unique properties, the molecular and chemical determinants underlying these biofilm-phenotypes remain poorly understood. Herein, we aim to identify and quantify the components in the biofilm ECM and how they modulate radical stability. Characterizing native biofilms at molecular resolution remains experimentally challenging because of their compositional complexity and structural heterogeneity. Only recently, we used solid-state NMR (ssNMR) to examine biofilm systems.^[4, 28-30]^ However, acquiring highly sensitive spectra typically requires high sample concentrations, advanced multidimensional methods, or isotopic labeling, which can be impractical for native biofilms, particularly those obtained from patient-derived samples. Dynamic nuclear polarization (DNP) ssNMR increases signal sensitivity by several orders of magnitude compared to conventional ssNMR, utilizing hyperpolarization from stable nitroxide radicals, known as polarization agents.^[31-39] [40]^ Recently, we extended DNP ssNMR to native *P. fluorescens* colony biofilms and observed a remarkably high DNP enhancement of ∼75 using the nitroxide-based polarization agent AsymPokPOK.^[40]^ However, we also observed a decay in DNP enhancement during the incubation of biofilm with AsymPolPOK at room temperature, indicating degradation of radical within the native biofilm environment.^[40]^ These observations indicate that biofilms contain chemically heterogenous environments capable of strongly modulating radical stability and small-molecule accessibility. The structural origins and location of this nitroxide radical decay in biofilm systems are important to decipher, as it can provide key insights into the function that the ECM plays in biofilm-mediated antimicrobial resistance.

In this work, we determine whether differences in biofilm ECM composition create distinct reductive environments that influence nitroxide stability within native biofilms. Using complementary electron paramagnetic resonance (EPR) and DNP ssNMR measurements, we quantify the impact of the Pf0-1 biofilm environment on radicals and correlate the effects with structural and compositional features of the ECM. In these measurements, EPR provides quantitative insights into radical degradation, while conventional and DNP ssNMR reveals complementary information about the ECM composition, organization, and molecular interactions with the radicals. By pairing these techniques, we gain a comprehensive understanding of small molecule diffusion in these native and complex biological environments and provide insight into nitroxide-based antibiofilm therapeutic strategies.

## Results and Discussion

### Reduction of Radicals by Biofilm is Dependent on ECM Composition

To investigate how different structural and chemical states of bacterial communities influence redox activity, we prepared three distinct *P. fluorescens* samples: wild-type (WT) biofilm, mucoid biofilm, and dry biofilm.^[27]^ The WT biofilm consists of surface-adhered bacterial cells embedded within a self-produced extracellular matrix (ECM). In contrast, the mucoid and dry biofilms were formed from previously isolated genetic variants of the WT strain and exhibit markedly different ECM features.^[41, 42]^

**Figure 1A** shows the electron microscopy (EM) micrographs and schematic illustrations of each sample, revealing that all three biofilms contain distinct abundant ECM, evident from the amorphous structures surrounding the cells in the micrographs. The WT biofilm reflects the native architecture formed under standard growth conditions. The mucoid variant is enriched with a glucose-rich polymer matrix, yielding a relatively loose and gel-like consistency.^[27, 41]^ Functionally, mucoid nature in *P. aeruginosa* biofilm is associated with an increased propensity of virulence of the bacteria.^[43]^ Despite its prevalence in upper respiratory infections, isolated mucoid *P. aeruginosa* has shown decreased resistance to antibiotics compared to non-mucoid morphologies.^[44, 45]^ Interestingly, while there is not a direct correlation between high mucoid morphology and antibiotic resistance, mixed communities of mucoid and non-mucoid biofilm show a marked increase in resistance to endogenous antimicrobials.^[46]^ In comparison, the dry biofilm exhibits a dense and wrinkled morphology typically associated with more rigid morphology and being less hydrated.^[27]^ The wrinkles in the dry phenotype provide an increased surface-to-volume ratio, which can promote flow of nutrients throughout the biofilm for survival.^[47-49]^ These compositional and morphological differences reflect distinct microenvironments within the biofilms. Given the variation in ECM structure and hydration levels, we hypothesized that each sample would impact the diffusion of small molecules throughout the biofilm, thereby presenting unique redox profiles and resistance.

**Figure 1.**
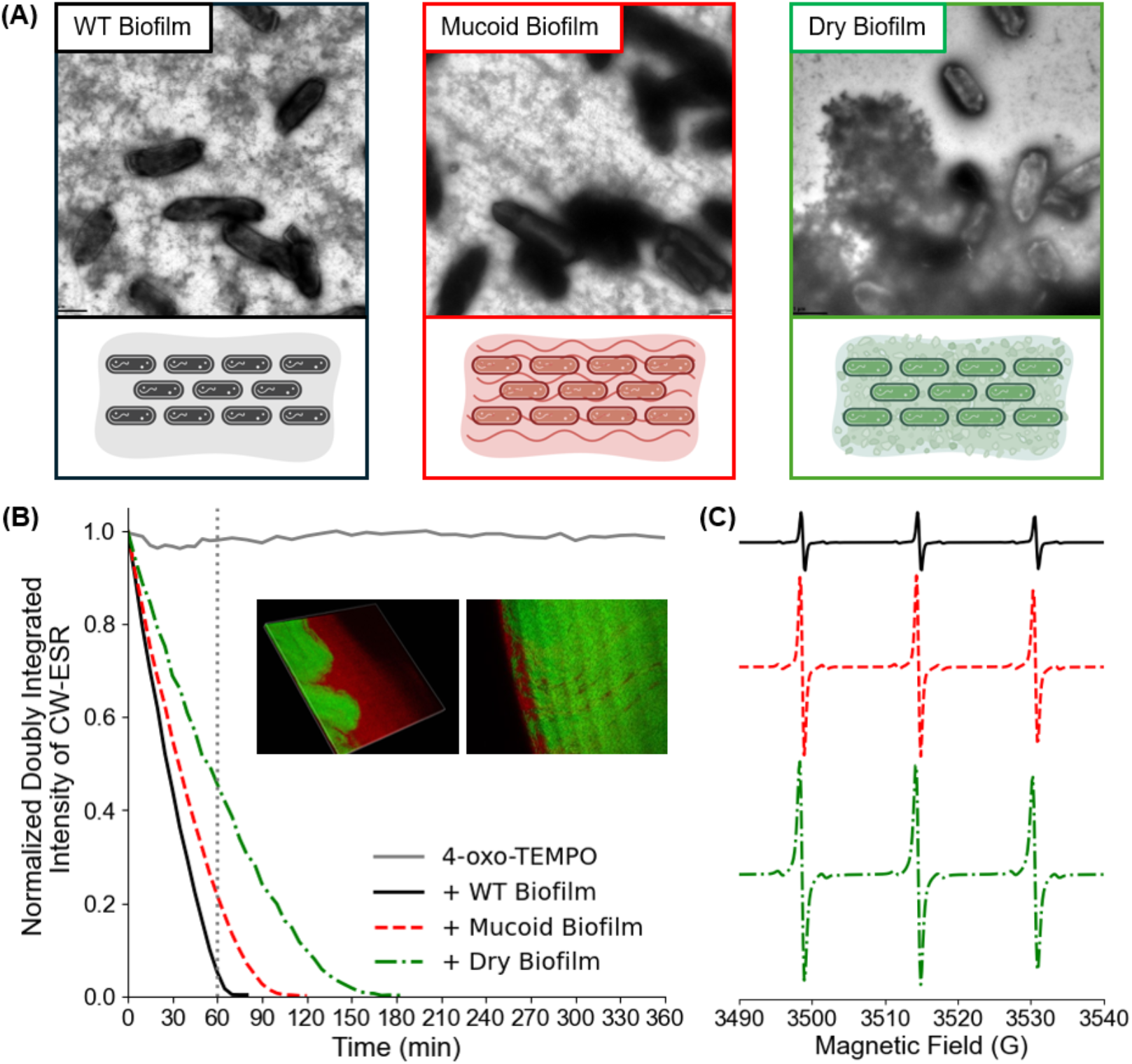
(A) Negative-stain EM micrographs of wild-type (WT) biofilm, mucoid biofilm, and dry biofilm. Each EM image is accompanied by a cartoon created with BioRender.com. (B) Doubly integrated intensity of continuous-wave (CW) EPR spectra as a function of time. Comparison of 4-oxo-TEMPO with the radical in WT biofilm (solid black), mucoid biofilm (dashed red), and dry biofilm (dash-dotted green). The inset shows the confocal laser scanning microscopy images of biofilms formed by co-cultures of the mucoid (red, tagged with RFP) and dry (green, tagged with GFP) variants as they auto segregate due to their ECM differences. (C) CW-EPR spectrum of each sample at the 60-minute time point, indicated by the dotted grey vertical line in Figure 1B.

To test this hypothesis, we initially employed the monoradical 4-oxo-TEMPO for subsequent EPR experiments. The use of a monoradical provides a straightforward measure of overall reductive potential while avoiding complications from intramolecular exchange interactions inherent in biradicals such as AsymPolPOK. Continuous-wave (CW) EPR spectroscopy was used to monitor the temporal decay of the 4-oxo-TEMPO signal upon incubation with the different bacterial samples. **Figure 1B** shows the normalized doubly integrated intensity of the spectrum over time, reflecting the total radical concentration in the system. Representative CW-EPR spectra at the 60-minute time point, shown in **Figure 1C**, illustrate differences in signal intensity across samples.

In WT biofilms, the EPR signal diminished to baseline within 70 minutes at room temperature. Both mucoid and dry biofilms also reduced the 4-oxo-TEMPO radical, albeit at slower rates than the intact WT biofilm. These differences are consistent with their distinct ECM compositions. In particular, the dry biofilm, characterized by its denser morphology and lower water content, exhibits the slowest reduction rate, reaching complete quenching of the EPR signal after approximately three hours.

Although mixed-culture reduction was not directly examined by EPR, prior observations of co-cultured variants provide useful context. When mucoid and dry variants are grown together, the dry biofilm forms atop a mucoid layer. Confocal laser scanning microscopy images, shown in **Figure 1B**, presents a continuous mucoid sheet surrounding the dry variant at the colony edge. This particular spatial segregation phenotype is unique to mixed cultures of mucoid and dry variants.^[27]^ In contrast, co-cultures of WT and mucoid variants display a different pattern, with mucoid cells forming localized patches throughout the WT population and away from the colony edge.^[42]^ These different spatial organizations in mixed colonies reflect underlying differences in ECM composition and regulation, which may also influence redox behavior.^[27, 41]^

In contrast to biofilms, planktonic cells are free-floating and lack an ECM, serving as the initial state prior to biofilm formation.^[50-52]^ To create an accurate comparison of planktonic cells to the densely packed environment of biofilm, we used pelleted planktonic cells to prepare our subsequent EPR samples. Under equivalent cell mass-to-radical volume ratios, established via cell counting (**Figure S1**), the CW-EPR signal for planktonic cells (**Figure 2A**) decayed significantly faster than the biofilm samples (**Figure 2B**). At a ratio of 10 mg of cells to 10 μL of TEMPO solution (1X), the planktonic cells completely reduced the EPR signal in just 15 minutes, while the equivalent biofilm sample took twice the amount of time to accomplish the same amount of reduction. Interestingly, the 2X (10 mg:20 μL) planktonic sample reduced the signal at an almost identical rate to the 1X biofilm sample. This trend continues through the rest of the data, with the 3X planktonic sample matching the 2X biofilm, and 5X planktonic matching the 3X biofilm.

**Figure 2.**
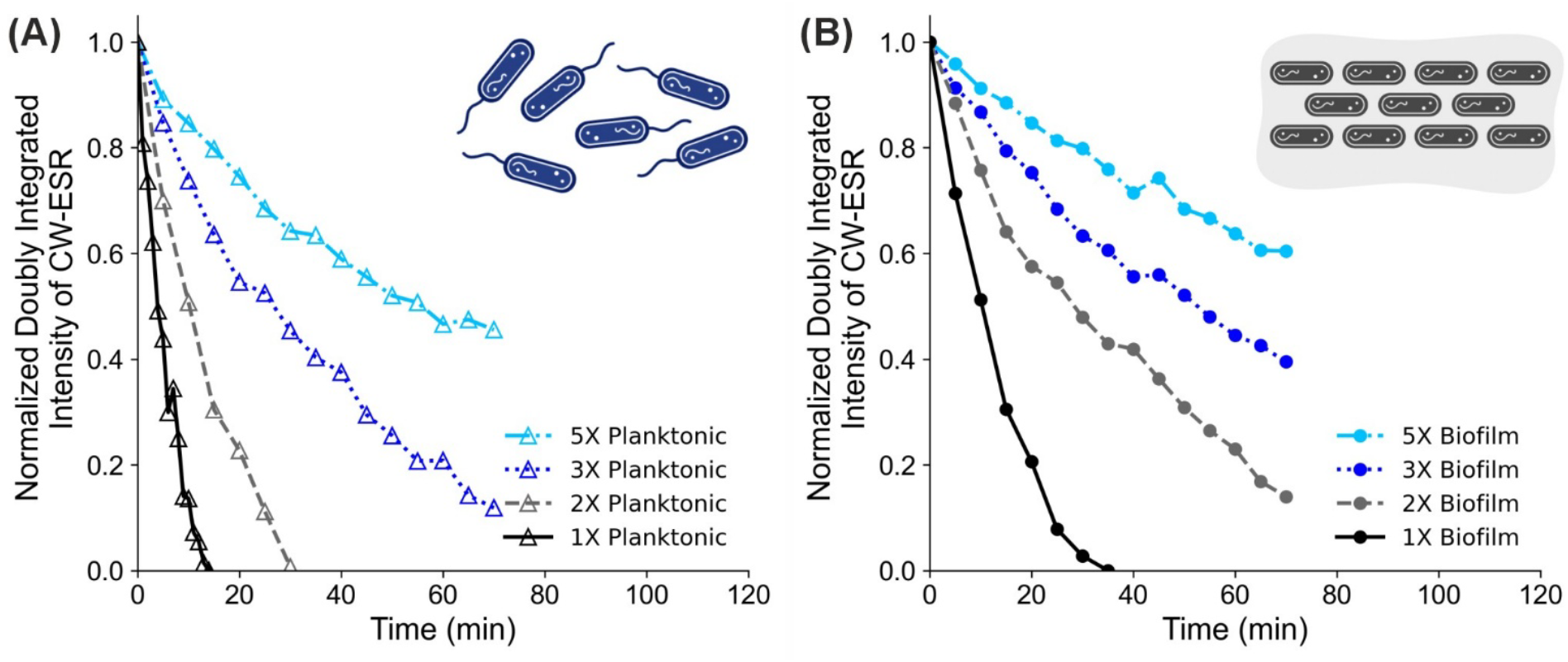
Normalized doubly integrated intensity of CW-EPR spectra as a function of time for (A) WT planktonic cells and (B) WT biofilm in the presence of 4-oxo-TEMPO. The insets show illustrations of planktonic cells (A) and WT biofilm (B) made using BioRender.com. The 1X samples (10 mg cells and 10 µL radical solution) are shown in black, 2X in grey, 3X in dark blue, and 5X in light blue.

These results suggest that the cells, rather than the ECM, are the primary reducing component within the biofilm system. While planktonic cells rapidly reduce the nitroxide signal, radical reduction occurs more slowly in intact biofilms, indicating that the ECM partially limits diffusion of the radical to the cellular reducing environment. One possible mechanism is similar to the response observed at low concentrations of nitric oxide,^[11]^ in which the nitroxide radical causes dispersion of the biofilm into free-floating planktonic cells that subsequently reduce the radical more efficiently.

To further probe this effect, a series of control experiments were conducted (see **Figure S2**). Extracted ECM from WT biofilm, which contains only the insoluble ECM fraction of the biofilm as demonstrated by our previously described method,^[28, 53]^ does not exhibit any radical reduction, indicating that the extracted ECM alone is not intrinsically redox-active. Nevertheless, it is important to note here that the chemical extraction protocol of ECM, may have removed/deactivated certain ECM components. Additionally, dehydrated WT biofilms (dried at 50 °C) showed no EPR signal decay, suggesting that increased ECM density or reduced hydration can effectively block radical penetration through the ECM. Finally, sonicated planktonic cells in dilute solution exhibited gradual signal decay, consistent with the release of intracellular reducing agents into solution.

Overall, the CW-EPR experiments with 4-oxo-TEMPO suggest that the ECM of biofilm functions as a protective barrier before the nitroxide radical potentially disperses the embedded cells, slowing the diffusion of small molecules into the reducing environment of the bacterial cells. Consequently, increased biomolecular density in the ECM, due to genetic changes in morphology or hydration levels, gives rise to distinct rates of small molecule diffusion across biofilm types. Once the embedded cells are dislodged from the ECM, the radical is easily reduced by the cellular environment.

### Stability of AsymPolPOK in Biofilm

Building on our results with 4-oxo-TEMPO, we next examined the stability of the AsymPolPOK polarization agent in biofilms. CW-EPR spectroscopy was used to monitor the redox state of AsymPolPOK over time, allowing us to assess its stability in complex biofilm and relevant environments. We compared how AsymPolPOK interacted with 1X WT biofilm and 1X planktonic samples. As shown in **Figure 3A**, both systems showed minimal reduction in total radical concentration over 180 minutes. Additionally, we show that in a dilute sample of WT biofilm and planktonic cells, the AsymPolPOK signal persists over 24 hours (see **Figure S3**). Remarkably, in comparison to mammalian cell lysates, for example, the biradical signal in these biofilm preparations persists 4-to 20-fold longer.^[54, 55]^

**Figure 3.**
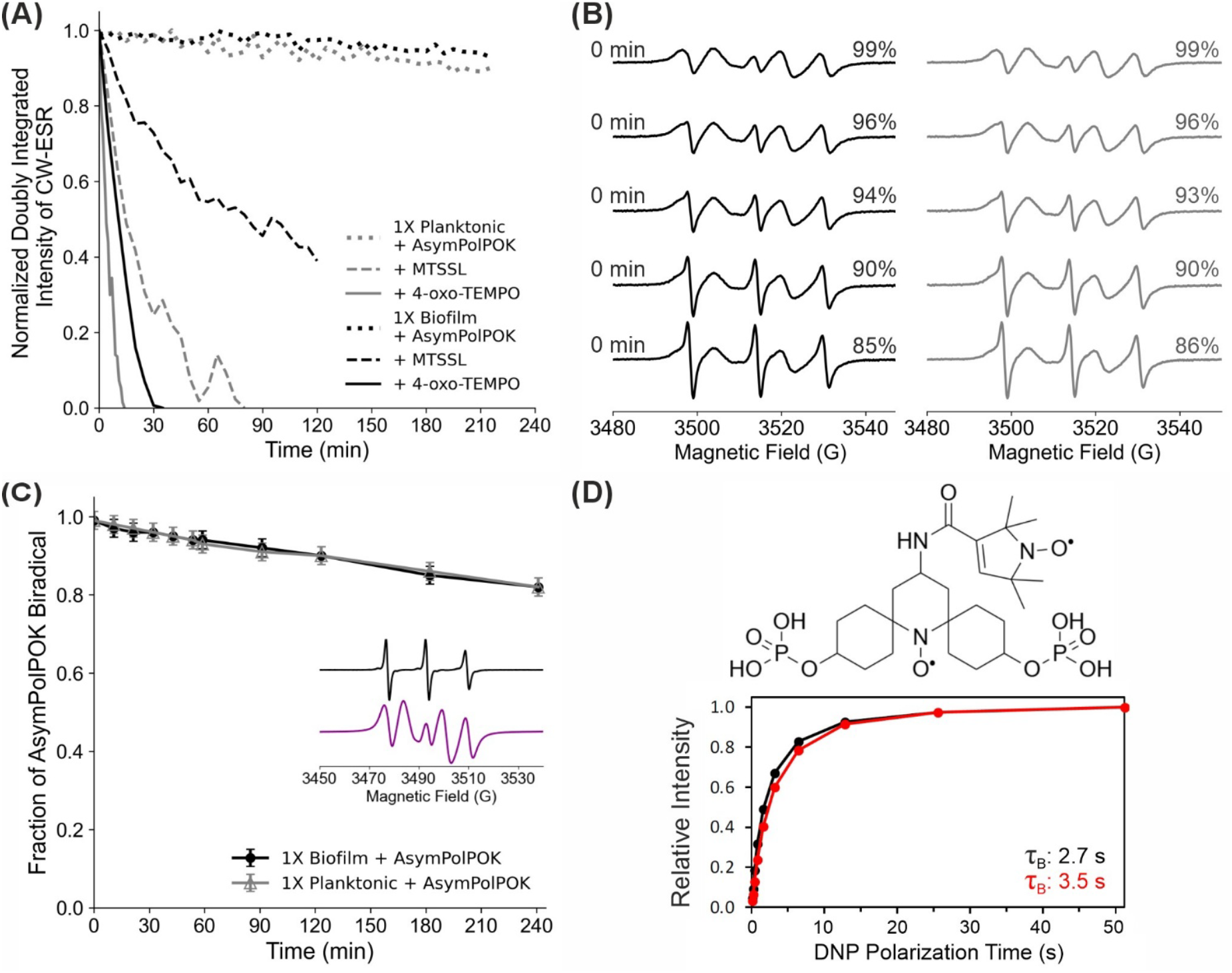
(A) Radical stability in WT biofilm and planktonic cells. Doubly integrated intensity of CW-EPR spectra as a function of time for 1X WT biofilm (black) and 1X planktonic cells (grey) with AsymPolPOK (dotted), MTSSL (dashed), and 4-oxo-TEMPO (solid). 1X TEMPO data is repeated from Fig. 2 for direct comparison. (B) CW-EPR spectra as a function of time for the AsymPolPOK data in Figure 3A: WT biofilm (left, black) and WT planktonic cells (right, grey). Time is increasing on the y-axis in 30-minute intervals. The spectra were simulated by adding DNP active and inactive AsymPolPOK in different ratios to achieve a good fit. The percentages shown are the fractions of DNP active AsymPolPOK in each spectrum as evaluated by the simulation. (C) Fraction of AsymPolPOK radical in the EPR spectra. The inset shows the two component spectra used for the simulations. (D) Chemical structure of DNP active AsymPolPOK. DNP buildup times (τ_B_) as a function of polarization time for active (black) and partially active (red) AsymPolPOK.

To gain a deeper understanding of the chemoselective reduction taking place in the cells, we also evaluated the reduction of monoradicals structurally related to the individual moieties in AsymPolPOK (**Figure S4**): 4-oxo-TEMPO and (1-oxyl-2,2,5,5-tetramethylpyrroline-3-methyl)methanethiosulfonate spin label (MTSSL). **Figure 3A** highlights the increased resistance of AsymPolPOK to reduction in biofilms compared to the monoradicals 4-oxo-TEMPO and MTSSL, consistent with its design as a more robust polarizing agent. Additionally, the data show the increased reduction resistance of five-membered rings over six-membered rings, in agreement with prior reports.^[54, 56-59]^

Although the total AsymPolPOK radical concentration remains largely unchanged over time, this does not prevent partial reduction of the polarization agent into its DNP inactive form. **Figure 3B** shows the CW-EPR spectra over time as AsymPolPOK is reduced. Initially, the spectrum displays five distinct lines, characteristic of strong exchange interactions between the two nitroxide moieties in the biradical.^[60, 61]^ However, as incubation progresses at room temperature, the spectral lines evolve into a more typical single nitroxide EPR spectrum, becoming dominated by the nitrogen hyperfine interaction, losing the five-line pattern associated with exchange coupling. To quantify this process, we simulated the CW-EPR spectra by combining the DNP inactive spectrum, obtained after incubating AsymPolPOK with biofilm for 48 hours, with the DNP active spectrum. (see **Figure S5**). **Figure 3C** illustrates the gradual decrease in the DNP active fraction of AsymPolPOK as the WT biofilm and planktonic cells reduce the radical. Further explanation of the simulation process can be found in the SI (Figure S5). Interestingly, although the total radical concentration remains relatively stable, slow reduction of one radical moiety does occur within the 3-hour period. The continued persistence of the triplet EPR signal, even at later time-points, indicates asymmetric protection of the two radical centers. By comparison of CW-EPR spectra of five- and six-membered nitroxides to the reduced AsymPolPOK spectrum, we suggest that the six-membered ring is selectively reduced by cellular environments (see **Figure S4**), which aligns with prior observations.^[54, 61, 62]^

These results demonstrate that, despite the relative stability in total radical concentration, slow conversion of AsymPolPOK to its DNP inactive form occurs on the hour(s) timescale. To evaluate how this redox transformation affects DNP efficiency, we recorded ^13^C CP MAS ssNMR spectra of WT biofilm at both early and late stages of incubation with AsymPolPOK. After prolonged incubation, there is a clear reduction in DNP signal enhancement (**Figure S6**) due to a decrease in the cross-effect upon the reduction of one nitroxide radical in AsymPolPOK. Measurements of DNP buildup times (τ_B_) at 100 K under microwave irradiation, as shown in **Figure 3D**, reveal a longer polarization buildup for the partially reduced AsymPolPOK sample, consistent with reduced polarization transfer efficiency due to reduction in biradical concentration. Notably, this represents the first demonstration of AsymPolPOK signal decay in biofilms which directly links the decay of one radical in the polarization agent to a decrease in DNP efficiency.

We also investigated whether oxidizing conditions can inhibit radical reduction in biofilms to increase DNP efficiency at later time-points. WT biofilms exposed to 4-oxo-TEMPO were treated with hydrogen peroxide (H_2_O_2_) or potassium ferricyanide (K_3_Fe(CN)_6_) (see **Figure S7**). Excessive concentrations of hydrogen peroxide are known to be cytotoxic, so we employed the more biocompatible potassium ferricyanide as an alternative solution.^[63-66]^ In both cases, the radical lifetime of 4-oxo-TEMPO increased significantly, by almost four-fold with hydrogen peroxide and more than eight-fold with potassium ferricyanide.

Collectively, these EPR results reveal that nitroxide radical behavior in biofilms is determined by cellular reducing activity, nitroxide-induced dispersion into planktonic cells, ECM-mediated diffusion, and the intrinsic stability of AsymPolPOK. These findings also highlight the importance of both the local biofilm microenvironment and polarization agent design to optimize DNP performance.

### High-resolution ssNMR Identifies Radical Interaction Sites in the Biofilm

While the EPR experiments suggest that planktonic cellular environments are primarily responsible for nitroxide reduction, the ECM plays a major role in slowing down radical reduction. Therefore, we next sought to determine how the radical interacts with the biofilm ECM to reduce radical penetration into planktonic cells. The ECM appears to function as a diffusion and transport barrier, so it is important to identify the biomolecular components that localize the radical within the matrix. To identify the specific molecules, we performed ssNMR measurements on WT biofilm samples in the presence and absence of 4-oxo-TEMPO.

Cross polarization (CP)- and INEPT-based ^13^C spectra were acquired at room temperature to probe rigid and mobile structural components of the biofilm sample, respectively. CP spectra report on structurally rigid components such as polysaccharides and lipids, while the INEPT spectra detect highly mobile and flexible components, including metabolites and dynamic ECM molecules, as we have shown previously.^[30, 67, 68]^ **Figure 4A** presents the ^13^C CP ssNMR spectra of the native WT biofilm, revealing characteristic signals from proteins, polysaccharides, nucleic acids, lipids, and other biofilm constituents, consistent with previous assignments.^[30, 40, 69]^ Upon addition of 4-oxo-TEMPO, a reduction in relative spectral intensity is observed across multiple regions. The largest decreases occur at ∼35 ppm and between 50-110 ppm, corresponding primarily to lipid and polysaccharide signals, respectively. Carbonyl (CO) signals are also slightly reduced upon radical addition. The signals between 10-25 ppm and 110-170 ppm were not affected by the radical and were therefore used as normalization reference points.

**Figure 4.**
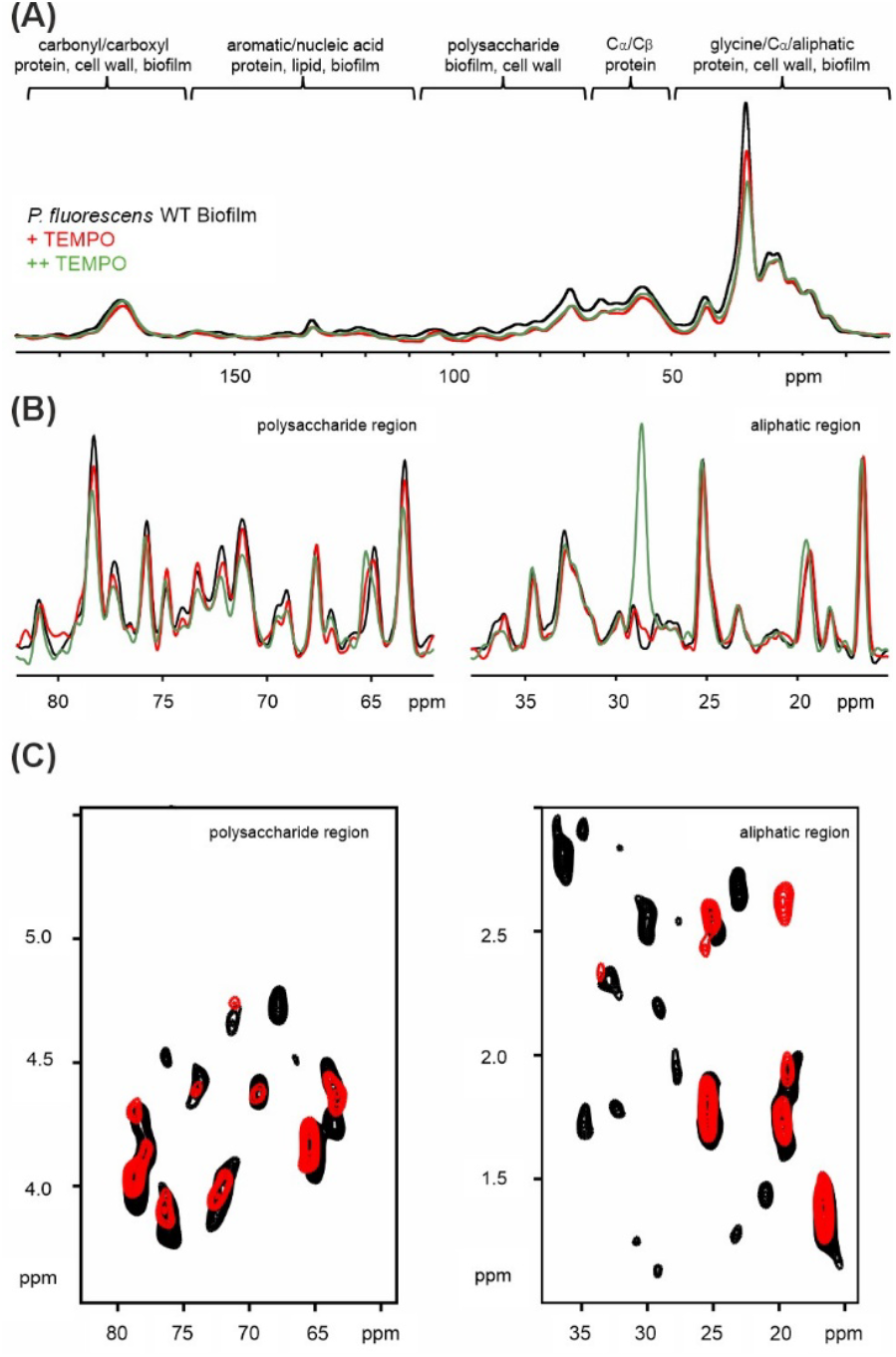
(A) 1D ^13^C CP conventional ssNMR spectrum of the WT bacterial biofilm (black) without TEMPO, and with 10 mM (red) and excess (green) TEMPO. (B) 1D ^13^C INEPT ssNMR spectra of the polysaccharide region (80 – 60 ppm) (left) and the aliphatic region (40 – 15 ppm) (right). The additional signal at ∼28 ppm is due to the excess added radical and represents an impurity signal. (C) 2D ^1^H-^13^C INEPT ssNMR spectra in the absence (black, ^1^H-detected) and presence (red, ^1^H-detected) of TEMPO. The artefact at ∼3 ppm in the red spectrum is due to the remaining signal from the TEMPO buffer that was not suppressed efficiently compared to the water suppression. As a reference and visual guide, tentative resonance assignments of the protein amino acid and polysaccharide signals in a different WT biofilm sample, recorded with more scans, is given (blue, ^13^C-detected), as reported previously.^[30]^ Approximate ^1^H/^13^C NMR resonance regions corresponding to polysaccharide and protein/aliphatic components are marked with dashed lines.

Overall, resonances observed between 30-110 ppm decreased by approximately 30%. This indicates that the radical interacts with lipids and polysaccharides, which may be related to the interaction interface and reduction mechanism of radical in biofilm. At a higher concentration of 4-oxo-TEMPO, the lipid signal at ∼35 ppm is further reduced by an additional 20%, suggesting a preferential interaction of the nitroxide radical with lipids. Excess radical did not reduce other signals. Protein resonances may also contribute to both the 50-65 ppm region and the CO region, so interactions between the radical and the protein components of the ECM or biofilm cannot be excluded by 1D ssNMR spectra and requires 2D ssNMR spectra as described below.

To further resolve the flexible biofilm components, ^13^C INEPT ssNMR spectra highlighting the polysaccharide and aliphatic carbon regions are shown in **Figure 4B**. Similar to the CP spectra, the aliphatic signals did not exhibit any reduction in intensity upon radical addition. On the other hand, the polysaccharide resonances exhibit a strong, concentration-dependent decrease, as we assigned those resonances previously by 2D ssNMR spectra as shown here in **Figure 4C**. Although individual polysaccharide peaks could not be assigned due to the complexity of the 1D spectra and overlap, the selective changes in this region indicate that 4-oxo-TEMPO interacts predominantly with the flexible polysaccharide components of the biofilm ECM, as also observed for the rigid fraction by CP.

Further confirmation for this conclusion comes from the 2D ^1^H-^13^C INEPT spectra acquired in the presence and absence of 4-oxo-TEMPO. **Figure 4C** shows that radical-mediated signal reduction predominantly occurs for the resonances which do not coincide with the protein resonances previously assigned in a WT *Pseudomonas* biofilm sample recorded with higher signal-to-noise (blue spectrum with tentative assignments). This confirms that the NMR signal intensity loss observed is due to interactions of 4-oxo-TEMPO and the polysaccharide-rich fraction of the biofilm rather than the protein components.

The CP and INEPT ssNMR spectra overall strongly indicate that 4-oxo-TEMPO interacts with the rigid and flexible polysaccharide and rigid lipid components of the ECM or biofilm. These results indicate that the ECM is not only a diffusion barrier, but also a primary interaction site that determines radical localization within the biofilm. The selectivity of the nitroxide association with these ECM components provides a molecular-level explanation for the diffusion-limited reduction behavior observed in the EPR experiments.

### Ultrasensitive DNP ssNMR Provides Insights into Chemical Composition Differences in Biofilm Strains

After establishing that 4-oxo-TEMPO preferentially associates with polysaccharide and lipid-rich regions of the ECM, we next applied DNP-enhanced ssNMR spectroscopy to evaluate how the interactions translate into ultrasensitive structural determinations across the different biofilm variants. Since nitroxide radicals are known to modulate biofilm dispersal and antimicrobial efficiency, differences in ECM composition between these variants may directly impact radical accessibility and reduction behavior within biofilms. We performed ultrasensitive DNP-enhanced ssNMR spectroscopy using the AsymPolPOK biradical incorporation protocol described in Methods, achieving DNP enhancements of e∼75.^[40]^

High-sensitivity 1D ^13^C CPMAS and 2D ^1^H-^13^C and ^13^C-^13^C ssNMR spectra were recorded for native WT, mucoid, and dry biofilm samples at natural isotopic abundance and without additional treatment, as shown in **Figure 5A**.^[40]^ As a proof of concept, we demonstrated the feasibility of recording such multidimensional ssNMR spectra on WT biofilm samples (**Figures 5A-C**),^[40]^ and here we extend this DNP ssNMR approach to planktonic cells and different biofilm strains with distinct ECM properties.

**Figure 5.**
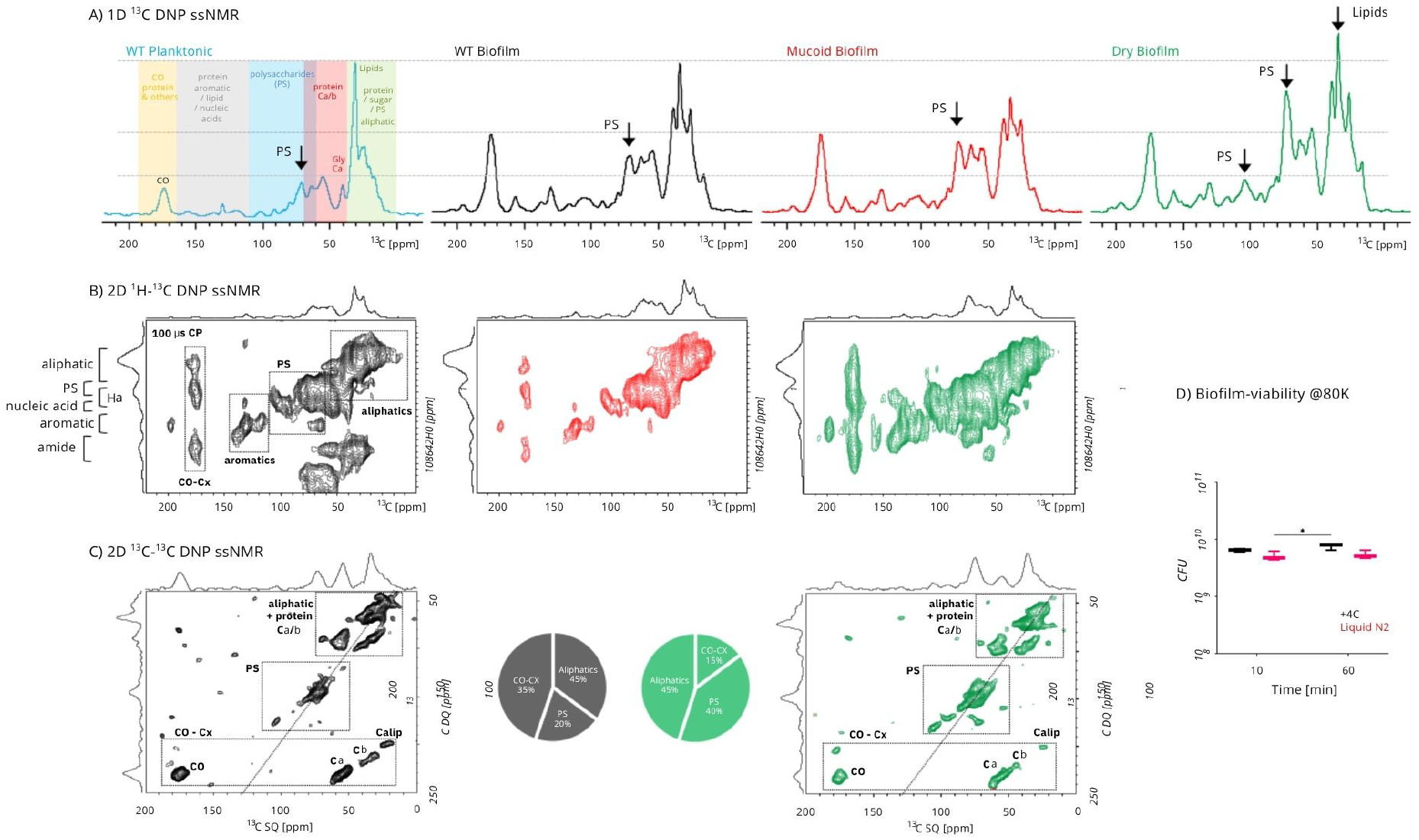
DNP ssNMR applications on *Pseudomonas* biofilms. (A) 1D ^13^C CPMAS DNP ssNMR spectra of WT planktonic (blue) and different biofilm strains: WT (black), Mucoid (red), and Dry (green) biofilm samples. (B) 2D ^1^H-^13^C CPMAS DNP ssNMR spectra of WT (black), Mucoid (green) and Dry (red) biofilm samples. 100 μs CP contact times were utilized. (C) 2D ^13^C-^13^C CPMAS refocused dipolar INADEQUATE SQ-DQ DNP ssNMR spectra of WT (black) and Dry (red) biofilm samples. The quantification of aliphatic, polysaccharide (PS), and carbonyl (CO) region cross peaks by integration of the given chemical shift ranges in the DQ F2 dimension for the Dry-variant biofilm sample is shown. The percentage abundances of corresponding chemical shifts are plotted in a pie-chart. The total experimental times are ∼5, ∼30, and ∼1000-1500 minutes for A, B and C, respectively. (D) The biofilm cell-viability assay represents the effect of keeping a fresh biofilm sample in the refrigerator (+4 °C) and in liquid nitrogen (∼80 K) for 10 and 60 minutes.

**Figure 6.**
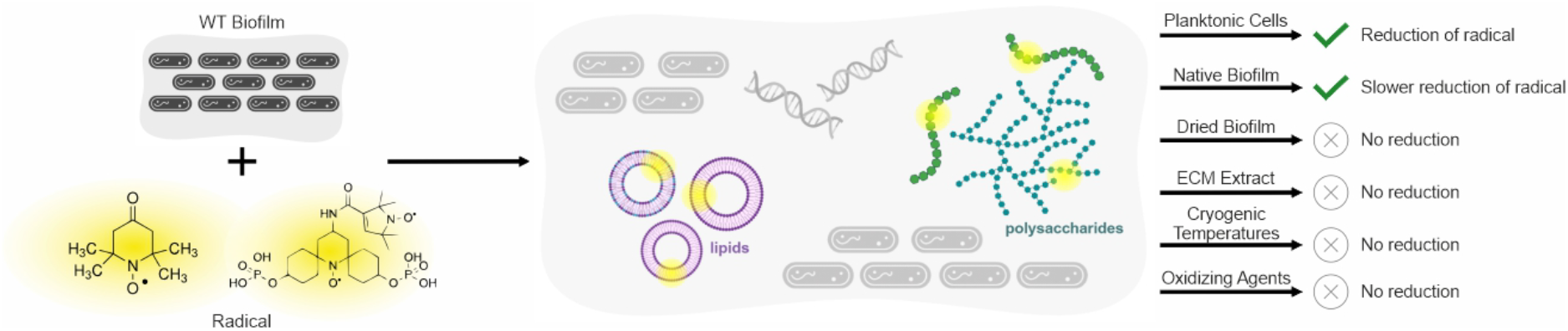
Summary of results. Upon addition of 4-oxo-TEMPO or AsymPolPOK to the native biofilm, the radical interacts primarily with lipids and polysaccharides in the biofilm ECM. There is a reduction of the radical by both planktonic cells and the WT biofilm when the radical interacts with the cell, with a slower reduction exhibited by the biofilm samples. On the other hand, dried WT biofilm, ECM extract, and biofilm under cryogenic temperatures do not reduce the radical. The addition of an oxidizing agent, such as hydrogen peroxide or potassium ferricyanide, slows the rate of reduction of the radical.

**Figure 5A** shows 1D ^13^C CPMAS DNP ssNMR spectra from WT planktonic cells, mucoid biofilm, and dry biofilm samples and compares to WT biofilm spectrum we previously demonstrated.^[40]^ The spectra were normalized either to the aliphatic signal at ∼35 ppm (for planktonic and WT biofilm samples) or to the CO signal at ∼176 ppm (for the mucoid and dry samples) to identify relative changes in composition. Distinct compositional trends were observed across the samples. As expected, relative polysaccharide signals increase from planktonic cells to WT biofilm. This increase continued in the mucoid variant, indicating that the secreted ECM polysaccharide species are the driving force for mucoid biofilm formation. From mucoid to dry biofilm, the relative abundance of the polysaccharide signals increased further, particularly in the ∼73 and ∼104 ppm regions, due to the increased abundance of specific polysaccharide types in the dry variant. These trends correlate well with the enhanced production of extracellular matrix predominantly composed of polysaccharides and proteins, as we quantified recently.^[27, 40]^

Moreover, the compositional trends correlate with their known biological differences. Specifically, the gradually increasing polysaccharides signatures constitute the WT-to-mucoid differences, which manifests more significantly in the mucoid-to-dry differences.^[27]^ These distinctions are shown in **Figures 5A** and **5B** as quantitative 1D ^13^C and 2D ^1^H-^13^C CPMAS correlation spectra, as previously demonstrated on a WT biofilm sample.^[40]^ Such changes likely influence radical partitioning and diffusion to cellular reducing environments, providing a structural basis for the radical interaction and thereby reduction behavior observed by EPR. To further resolve chemical compositional differences without spectral overlap, 2D ^13^C-^13^C CPMAS refocused dipolar INADEQUATE SQ-DQ DNP ssNMR spectra were acquired for the dry biofilm sample and compared to WT biofilm spectra as we previously showed,^[40]^ (**Figure 5C**). These experiments enabled direct and ultrafast analysis and quantification of different biofilm chemical components. The spectra reveal an approximately two-fold increase in relative polysaccharide content, rising from 20% in WT to 40% in dry samples, which correlates with the stiffened morphology of the dry variant.^[27]^

To confirm that liquid nitrogen temperatures utilized in DNP ssNMR experiments do not perturb the biofilms, we performed biofilm viability assays and compared biofilm samples stored at ∼4 °C and ∼80 K. As shown in **Figure 5D** there was no observable cell death detected upon freezing the biofilms, consistent with observations in mammalian cells.^[70]^

Across all strains, DNP ssNMR reveals that biofilm composition evolves in a systematic manner, with a progressive increase in polysaccharide-rich domains from WT to mucoid to dry variants. Together with the EPR measurements, these findings support a model in which ECM composition regulates nitroxide radical localization and access to cellular reducing environments, potentially influencing nitroxide-mediated biofilm disruption processes.

## Conclusion

In this work, we presented an integrative magnetic resonance study and combined EPR, conventional ssNMR, and DNP-enhanced ssNMR spectroscopy to determine how native *P. fluorescens* Pf0-1 colony biofilms regulate the accessibility, localization, and stability of nitroxide-based radicals, including 4-oxo-TEMPO and the DNP polarizing agent AsymPolPOK. Across all techniques, we show that biofilms actively regulate radical behavior through a combination of ECM-mediated diffusion barriers and selective molecular interactions within the ECM.

Biofilms protect bacterial communities through a variety of structural and biochemical mechanisms that collectively create a resilient microenvironment. The ECM limits the diffusion of harmful agents and forms a skin-like layer that protects hydration.^[71]^ Structural components such as amyloid proteins and cellulose contribute to desiccation tolerance, while extracellular enzymes can sorb and metabolize toxins.^[71-74]^ Beyond these established protective mechanisms, our findings suggest that the polysaccharides and lipids in the ECM strongly influence the transport and diffusion of redox-active small molecules within native biofilms. Rather than an intrinsically redox-active component, the ECM appears to regulate radical accessibility to the highly reducing intracellular and planktonic cellular environments responsible for nitroxide reduction. EPR measurements establish that radical reduction is strongly dependent on biofilm morphology, hydration, and density. Native WT biofilms exhibit the fastest radical decay, while the glucose-rich mucoid variant displays intermediate behavior, and the dry variant shows the slowest reduction kinetics. Additionally, isolated ECM and manually dehydrated WT biofilm exhibit no radical reduction, indicating that either the absence of cells or restricted radical diffusion through dense ECM environments inhibits reduction. In contrast, planktonic cells and their lysates reduce nitroxide radicals efficiently, supporting the conclusion that intracellular or cellular reducing environments are the primary source of radical decay. These EPR results support the idea that the ECM slows radical transport into reducing cellular conditions rather than directly mediating the reduction itself.

Complementary high-resolution ssNMR and DNP-enhanced ssNMR identify the molecular origin of this behavior by pinpointing the primary biomolecules that interact with water-soluble small molecules in native biofilms. Polysaccharides and lipids found in the ECM dominate these interactions, proving that the ECM functions as a highly complex protective barrier to the bacteria cells within biofilms. DNP ssNMR further demonstrates that these interactions evolve across biofilm morphologies, with increasing polysaccharide-rich ECM domains correlating with slower radical reduction observed by EPR.

Importantly, nitroxide radicals and nitric oxide mimics are known to promote biofilm dispersal and the formation of planktonic cell states. In this context, our results suggest a mechanism in which nitroxides initially partition into polysaccharide-rich regions of the ECM before gradually accessing cellular reducing conditions. Once the radicals reach planktonic cells or intracellular environments, rapid reduction occurs by exposure to cellular reducing agents. Our results suggest that native biofilms regulate nitroxides through both diffusion barriers and dynamic ECM interactions within heterogeneous biofilm environments.

Overall, this work establishes a molecular-level understanding of how native biofilm ECM composition and morphology regulate the behavior of redox-active small molecules. We provide insight into how biofilm structure can influence antimicrobial and nitroxide-based therapeutic accessibility. We also demonstrate the unique nature of combined EPR and DNP-enhanced ssNMR approaches for probing chemical interactions in complex native biofilms.

## Supporting Information

Experimental details and Figures S1-S12 are provided within the Supporting Information. The authors have cited additional references within the Supporting Information.^[63-66, 75-78]^

## Acknowledgements

UA acknowledges support from the Department of Structural Biology, University of Pittsburgh School of Medicine (UPSOM) for access to the high-field NMR facility. UA acknowledges start-up funding by the UPSOM. UA acknowledges a Competitive Medical Research Fund grant by the UPMC Health System. WK was supported by funding from the National Institute of General Medical Sciences of the NIH 1R15GM132856. The National High Magnetic Field laboratory (NHMFL) is funded by the National Science Foundation Division of Materials Research (DMR-1644779 and DMR-2128556) and the State of Florida. A portion of this work was supported by the NIH P41 GM122698 and RM1-GM148766. FJS acknowledges support from a postdoctoral scholar award from the Provost’s Office at Florida State University. SS was funded by NSF BSF MCB 2407706. Cartoon Figures were created in Biorender.com.

## Entry for the Table of Contents

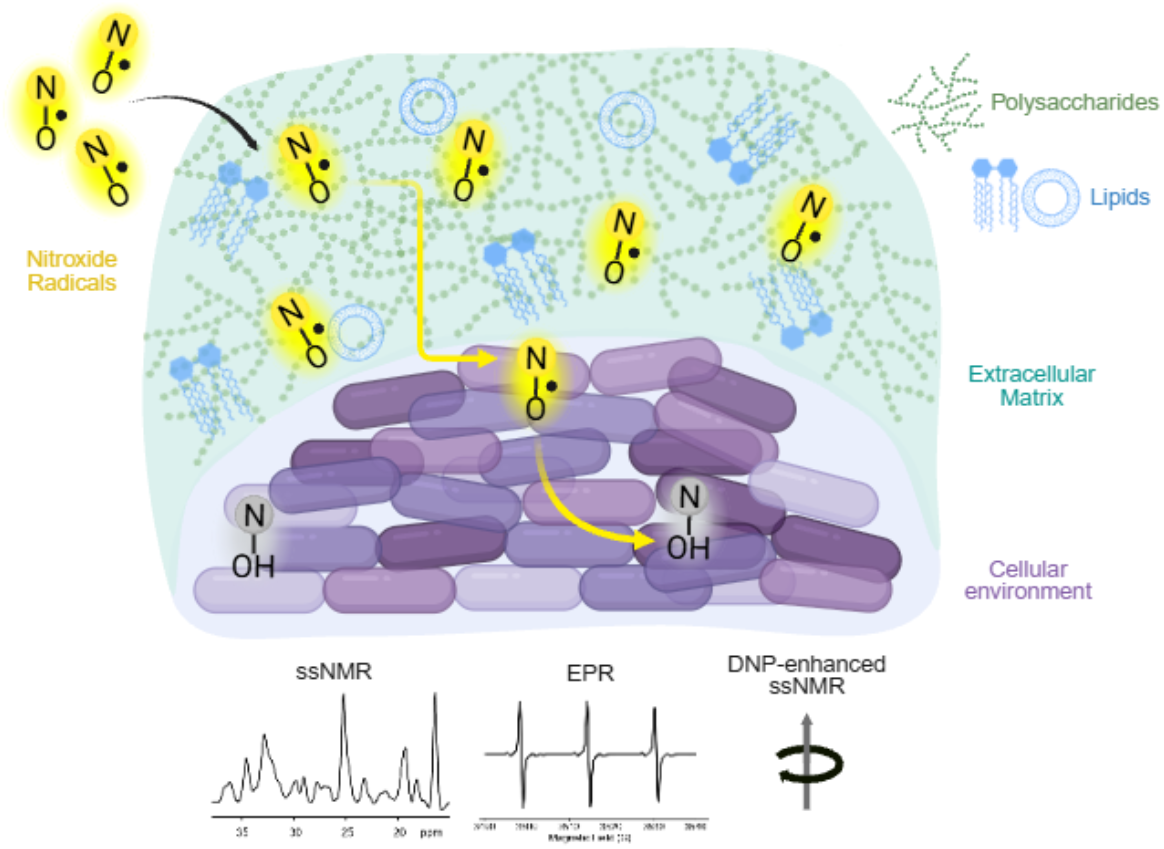

The complex biofilm environment consists of planktonic bacterial cells and cells embedded in an extracellular matrix (ECM). Through a combined magnetic resonance approach, we show that nitroxide radicals primarily interact with the polysaccharides and lipids in the ECM. Bacterial cells are responsible for the subsequent reduction of the radical. Created in BioRender.

## References

[1] Hoiby, N., Bjarnsholt, T., Givskov, M., Molin, S., Ciofu, O., “Antibiotic resistance of bacterial biofilms,” International Journal of Antimicrobial Agents 35, no. 4 (2010), 10.1016/j.ijantimicag.2009.12.011.

[2] Davies, D., “Understanding biofilm resistance to antibacterial agents,” Nature Reviews Drug Discovery 2(2003), 10.1038/nrd1008.

[3] Harimawan, A., Ting, Y.-P., “Investigation of extracellular polymeric substances (EPS) properties of P. aeruginosa and B. subtilis and their role in bacterial adhesion,” Colloids and Surfaces B: Biointerfaces 146(2016), 10.1016/j.colsurfb.2016.06.039.

[4] Akbey, Ü., Andreasen, M., “Functional amyloids from bacterial biofilms – structural properties and interaction partners,” Chem. Sci. 13, no. 22 (2022), 10.1039/D2SC00645F.

[5] Perry, E. K., Tan, M.-W., “Bacterial biofilms in the human body: prevalence and impacts on health and disease,” Front. Cell. Infect. Microbiol. 13(2023), 10.3389/fcimb.2023.1237164.

[6] Gebbink, M. F. B. G., Claessen, D., Bouma, B., Dijkhuizen, L., Wösten, H. A. B., “Amyloids — a functional coat for microorganisms,” Nat Rev Microbiol 3, no. 4 (2005), 10.1038/nrmicro1127.

[7] Collignon, P. J., Conly, J. M., Andremont, A., et al., “World Health Organization Ranking of Antimicrobials According to Their Importance in Human Medicine: A Critical Step for Developing Risk Management Strategies to Control Antimicrobial Resistance From Food Animal Production,” Clin Infect Dis 63, no. 8 (2016), 10.1093/cid/ciw475.

[8] Shaw, E., Wuest, W. M., “Virulence attenuating combination therapy: a potential multi-target synergy approach to treat Pseudomonas aeruginosa infections in cystic fibrosis patients,” RSC Med. Chem. 11, no. 3 (2020), 10.1039/C9MD00566H.

[9] Dieltjens, L., Appermans, K., Lissens, M., et al., “Inhibiting bacterial cooperation is an evolutionarily robust anti-biofilm strategy,” Nature Communications 11, no. 1 (2020): 107, 10.1038/s41467-019-13660-x.

[10] Flemming, H.-C., van Hullebusch, E. D., Neu, T. R., et al., “The biofilm matrix: multitasking in a shared space,” Nat Rev Microbiol 21, no. 2 (2023), 10.1038/s41579-022-00791-0.

[11] Barraud, N., Hassett, D. J., Hwang, S.-H., Rice, S. A., Kjelleberg, S., Webb, J. S., “Involvement of Nitric Oxide in Biofilm Dispersal of Pseudomonas aeruginosa,” Journal of Bacteriology 188, no. 21 (2006), 10.1128/jb.00779-06.

[12] Roberts, J. M., Milo, S., Metcalf, D. G., “Harnessing the Power of Our Immune System: The Antimicrobial and Antibiofilm Properties of Nitric Oxide,” Microorganisms 12, no. 12 (2024).

[13] de la Fuente-Nunez, C., Reffuveille, F., Fairfull-Smith, K. E., Hancock, R. E., “Effect of nitroxides on swarming motility and biofilm formation, multicellular behaviors in Pseudomonas aeruginosa,” Antimicrob Agents Chemother 57, no. 10 (2013), 10.1128/AAC.01381-13.

[14] Barraud, N., Storey, M. V., Moore, Z. P., Webb, J. S., Rice, S. A., Kjelleberg, S., “Nitric oxide-mediated dispersal in single-and multi-species biofilms of clinically and industrially relevant microorganisms,” Microb Biotechnol 2, no. 3 (2009), 10.1111/j.1751-7915.2009.00098.x.

[15] Cutruzzola, F., Frankenberg-Dinkel, N., “Origin and Impact of Nitric Oxide in Pseudomonas aeruginosa Biofilms,” J Bacteriol 198, no. 1 (2016), 10.1128/JB.00371-15.

[16] Webster, C. M., Shepherd, M., “The nitric oxide paradox: antimicrobial and inhibitor of antibiotic efficacy,” Emerg Top Life Sci 8, no. 1 (2024), 10.1042/ETLS20230114.

[17] Schmidt, I., Steenbakkers, P. J., op den Camp, H. J., Schmidt, K., Jetten, M. S., “Physiologic and proteomic evidence for a role of nitric oxide in biofilm formation by Nitrosomonas europaea and other ammonia oxidizers,” J Bacteriol 186, no. 9 (2004), 10.1128/JB.186.9.2781-2788.2004.

[18] Verderosa, A. D., Mansour, S. C., de la Fuente-Nunez, C., Hancock, R. E., Fairfull-Smith, K. E., “Synthesis and Evaluation of Ciprofloxacin-Nitroxide Conjugates as Anti-Biofilm Agents,” Molecules 21, no. 7 (2016), 10.3390/molecules21070841.

[19] Alexander, S. A., Kyi, C., Schiesser, C. H., “Nitroxides as anti-biofilm compounds for the treatment of Pseudomonas aeruginosa and mixed-culture biofilms,” Org Biomol Chem 13, no. 16 (2015), 10.1039/c5ob00284b.

[20] Verderosa, A. D., Hawas, S., Harris, J., Totsika, M., Fairfull-Smith, K. E., “Isothiazolone-Nitroxide Hybrids with Activity against Antibiotic-Resistant Staphylococcus aureus Biofilms,” ACS Omega 7, no. 6 (2022), 10.1021/acsomega.1c06433.

[21] verderosa, A. D., Dhouib, R. F.-S. K. E., Totsika, M., “Nitroxide Functionalized Antibiotics Are Promising Eradication Agents against Staphylococcus aureus Biofilms,” Antimicrobial Agents and Chemotherapy 64, no. 1 (2019), 10.1128/aac.01685-19.

[22] Michl, T. D., Tran, D. T. T., Kuckling, H. F., et al., “It takes two for chronic wounds to heal: dispersing bacterial biofilm and modulating inflammation with dual action plasma coatings,” RSC Adv 10, no. 13 (2020), 10.1039/c9ra09875e.

[23] Tan, X., Hu, M., Cheng, X., Xiao, J., Zhou, J., Zhu, G., “Effects of elevated levels of intracellular nitric oxide on Pseudomonas aeruginosa biofilm in chronic skin wound and slow-killing infection models,” Int Microbiol 27, no. 2 (2024), 10.1007/s10123-023-00395-5.

[24] Firoved, A. M., Wood, S. R., Ornatowski, W., Deretic, V., Timmins, G. S., “Microarray analysis and functional characterization of the nitrosative stress response in nonmucoid and mucoid Pseudomonas aeruginosa,” J Bacteriol 186, no. 12 (2004), 10.1128/JB.186.12.4046-4050.2004.

[25] Carpenter, A. W., Schoenfisch, M. H., “Nitric oxide release: part II. Therapeutic applications,” Chem Soc Rev 41, no. 10 (2012), 10.1039/c2cs15273h.

[26] Lewandowski, M., Gwozdzinski, K., “Nitroxides as Antioxidants and Anticancer Drugs,” Int J Mol Sci 18, no. 11 (2017), 10.3390/ijms18112490.

[27] Kim, W., Levy, S. B., Foster, K. R., “Rapid radiation in bacteria leads to a division of labour,” Nat Commun 7, no. 1 (2016), 10.1038/ncomms10508.

[28] Byeon, C.-H., Hansen, K. H., DePas, W., Akbey, Ü., “High-resolution 2D Solid-State NMR provides insights into Nontuberculous Mycobacteria,” (2024), 10.1101/2024.05.28.596255.

[29] Hansen, K. H., Golcuk, M., Byeon, C. H., et al., “Structural Basis of Pseudomonas FapC Biofilm-Forming Functional Amyloid Formation,” (2025), 10.1101/2025.03.15.642095.

[30] Byeon, C.-H., Kinney, T., Saricayir, H., et al., “Tapping into the native Pseudomonas bacterial biofilm structure by high-resolution multidimensional solid-state NMR,” Journal of Magnetic Resonance 357(2023), 10.1016/j.jmr.2023.107587.

[31] Frederick, K. K., Michaelis, V. K., Corzilius, B., et al., “Sensitivity-Enhanced NMR Reveals Alterations in Protein Structure by Cellular Milieus,” Cell 163, no. 3 (2015), 10.1016/j.cell.2015.09.024.

[32] Akbey, Ü., Franks, W. T., Linden, A., Orwick-Rydmark, M., Lange, S., Oschkinat, H. 2013, “Dynamic Nuclear Polarization Enhanced NMR in the Solid-State.” in Hyperpolarization Methods in NMR Spectroscopy. edited by Kuhn, L. T., Berlin, Heidelberg: Springer. ISBN 978-3-642-39728-8.

[33] Akbey, Ü., Oschkinat, H., “Structural biology applications of solid state MAS DNP NMR,” J Magn Reson 269(2016), 10.1016/j.jmr.2016.04.003.

[34] Chow, W. Y., De Paëpe, G., Hediger, S., “Biomolecular and Biological Applications of Solid-State NMR with Dynamic Nuclear Polarization Enhancement,” Chem. Rev. 122, no. 10 (2022), 10.1021/acs.chemrev.1c01043.

[35] Gauto, D., Dakhlaoui, O., Marin-Montesinos, I., Hediger, S., Paëpe, G. D., “Targeted DNP for biomolecular solid-state NMR,” Chem. Sci. 12, no. 18 (2021), 10.1039/D0SC06959K.

[36] Jaudzems, K., Bertarello, A., Chaudhari, S. R., et al., “Dynamic Nuclear Polarization-Enhanced Biomolecular NMR Spectroscopy at High Magnetic Field with Fast Magic-Angle Spinning,” Angewandte Chemie International Edition 57, no. 25 (2018), 10.1002/anie.201801016.

[37] Linden, A. H., Lange, S., Franks, W. T., et al., “Neurotoxin II Bound to Acetylcholine Receptors in Native Membranes Studied by Dynamic Nuclear Polarization NMR,” J. Am. Chem. Soc. 133, no. 48 (2011), 10.1021/ja206999c.

[38] Narasimhan, S., Scherpe, S., Lucini Paioni, A., et al., “DNP-Supported Solid-State NMR Spectroscopy of Proteins Inside Mammalian Cells,” Angewandte Chemie International Edition 58, no. 37 (2019), 10.1002/anie.201903246.

[39] Ni, Q. Z., Daviso, E., Can, T. V., et al., “High Frequency Dynamic Nuclear Polarization,” Acc. Chem. Res. 46, no. 9 (2013), 10.1021/ar300348n.

[40] Byeon, C.-H., Kinney, T., Saricayir, H., et al., “Ultrasensitive Characterization of Native Bacterial Biofilms via Dynamic Nuclear Polarization-Enhanced Solid-State NMR,” Angewandte Chemie (2025), 10.1002/ange.202418146.

[41] Evans, A. F., Wells, M. K., Denk, J., et al., “Spatial Structure Formation by RsmE-Regulated Extracellular Secretions in Pseudomonas fluorescens Pf0-1,” Journal of Bacteriology 204, no. 10 (2022), 10.1128/jb.00285-22.

[42] Kim, W., Racimo, F., Schluter, J., Levy, S. B., Foster, K. R., “Importance of positioning for microbial evolution,” Proceedings of the National Academy of Sciences 111, no. 16 (2014), 10.1073/pnas.1323632111.

[43] Kirov, S. M., Webb, J. S., O’May, C. Y., et al., “Biofilm differentiation and dispersal in mucoid Pseudomonas aeruginosa isolates from patients with cystic fibrosis,” Microbiology 153, no. Pt 10 (2007), 10.1099/mic.0.2007/009092-0.

[44] Wang, Y. W. A., Qu, J., Ma, R., Kang, W., Liu, Y., Xu, Y., “Comparison of in vitro antimicrobial susceptibility between mucoid and non-mucoid Pseudomonas aeruginosa and its guiding value for antibiotic therapy,” Microb Spectrum 13, no. 13 (2025), 10.1128/spectrum.00287-25.

[45] Owlia, P., Nosrati, R., Alaghehbandan, R., Lari, A. R., “Antimicrobial susceptibility differences among mucoid and non-mucoid Pseudomonas aeruginosa isolates,” GMS Hygiene and Infection Control 9, no. 2 (2014).

[46] Malhotra, S., Limoli, D. H., English, A. E., Parsek, M. R., Wozniak, D. J., “Mixed Communities of Mucoid and Nonmucoid Pseudomonas aeruginosa Exhibit Enhanced Resistance to Host Antimicrobials,” mBio 9, no. 2 (2018), 10.1128/mbio.00275-18.

[47] Jo, J., Price-Whelan, A., Dietrich, L. E. P., “Gradients and consequences of heterogeneity in biofilms,” Nat Rev Microbiol 20, no. 10 (2022), 10.1038/s41579-022-00692-2.

[48] Zhang, C., Li, B., Tang, J. Y., Wang, X. L., Qin, Z., Feng, X. Q., “Experimental and theoretical studies on the morphogenesis of bacterial biofilms,” Soft Matter 13, no. 40 (2017), 10.1039/c7sm01593c.

[49] Wilking, J. N., Zaburdaev, V., De Volder, M., Losick, R., Brenner, M. P., Weitz, D. A., “Liquid transport facilitated by channels in Bacillus subtilis biofilms,” Proc Natl Acad Sci U S A 110, no. 3 (2013), 10.1073/pnas.1216376110.

[50] Berlanga, M., Guerrero, R., “Living together in biofilms: the microbial cell factory and its biotechnological implications,” Microbial Cell Factories 15, no. 1 (2016), 10.1186/s12934-016-0569-5.

[51] Zhao, A., Sun, J., Liu, Y., “Understanding bacterial biofilms: From definition to treatment strategies,” Front. Cell. Infect. Microbiol. 13(2023), 10.3389/fcimb.2023.1137947.

[52] Flemming, H.-C., Wingender, J., “The biofilm matrix,” Nat Rev Microbiol 8, no. 9 (2010), 10.1038/nrmicro2415.

[53] McCrate, O. A., Zhou, X., Reichhardt, C., Cegelski, L., “Sum of the Parts: Composition and Architecture of the Bacterial Extracellular Matrix,” Journal of Molecular Biology 425, no. 22 (2013), 10.1016/j.jmb.2013.06.022.

[54] Ackermann, B. E., Lim, B. J., Elathram, N., Narayanan, S., Debelouchina, G. T., “A Comparative Study of Nitroxide-Based Biradicals for Dynamic Nuclear Polarization in Cellular Environments,” ChemBioChem 23, no. 24 (2022), 10.1002/cbic.202200577.

[55] Lagasca, D., Ghosh, R., Xiao, Y., Frederick, K. K., “Stability of the polarization agent AsymPolPOK in intact and lysed mammalian cells,” Journal of Magnetic Resonance 374(2025), 10.1016/j.jmr.2025.107864.

[56] Keana, J. F. W., Pou, S., Rosen, G. M., “Nitroxides as potential contrast enhancing agents for MRI application: Influence of structure on the rate of reduction by rat hepatocytes, whole liver homogenate, subcellular fractions, and ascorbate,” Magnetic Resonance in Medicine 5, no. 6 (1987), 10.1002/mrm.1910050603.

[57] Jagtap, A. P., Krstic, I., Kunjir, N. C., Hänsel, R., Prisner, T. F., Sigurdsson, S. T., “Sterically shielded spin labels for in-cell EPR spectroscopy: Analysis of stability in reducing environment,” Free Radical Research 49, no. 1 (2015), 10.3109/10715762.2014.979409.

[58] Karthikeyan, G., Bonucci, A., Casano, G., et al., “A Bioresistant Nitroxide Spin Label for In-Cell EPR Spectroscopy: In Vitro and In Oocytes Protein Structural Dynamics Studies,” Angewandte Chemie International Edition 57, no. 5 (2018), 10.1002/anie.201710184.

[59] Swartz, H. M., Sentjurc, M., Morse, P. D., “Cellular metabolism of water-soluble nitroxides: Effect on rate of reduction of cell/nitroxide ratio, oxygen concentrations and permeability of nitroxides,” Biochimica et Biophysica Acta (BBA) - Molecular Cell Research 888, no. 1 (1986), 10.1016/0167-4889(86)90073-X.

[60] Eaton, S. S., Woodcock, L. B., Eaton, G. R., “Continuous wave electron paramagnetic resonance of nitroxide biradicals in fluid solution,” Concepts in Magnetic Resonance Part A 47A, no. 2 (2018), 10.1002/cmr.a.21426.

[61] Stevanato, G., Casano, G., Kubicki, D. J., et al., “Open and Closed Radicals: Local Geometry around Unpaired Electrons Governs Magic-Angle Spinning Dynamic Nuclear Polarization Performance,” J. Am. Chem. Soc. 142, no. 39 (2020), 10.1021/jacs.0c04911.

[62] Tagami, K., Equbal, A., Kaminker, I., Kirtman, B., Han, S., “Biradical rotamer states tune electron J coupling and MAS dynamic nuclear polarization enhancement,” Solid State Nuclear Magnetic Resonance 101(2019), 10.1016/j.ssnmr.2019.04.002.

[63] Wipf, P., Xiao, J., Jiang, J., et al., “Mitochondrial Targeting of Selective Electron Scavengers: Synthesis and Biological Analysis of Hemigramicidin−TEMPO Conjugates,” J. Am. Chem. Soc. 127, no. 36 (2005), 10.1021/ja053679l.

[64] Lawless, M. J., Shimshi, A., Cunningham, T. F., Kinde, M. N., Tang, P., Saxena, S., “Analysis of Nitroxide-Based Distance Measurements in Cell Extracts and in Cells by Pulsed ESR Spectroscopy,” ChemPhysChem 18, no. 12 (2017), 10.1002/cphc.201700115.

[65] Singewald, K., Lawless, M. J., Saxena, S., “Increasing nitroxide lifetime in cells to enable in-cell protein structure and dynamics measurements by electron spin resonance spectroscopy,” Journal of Magnetic Resonance 299(2019), 10.1016/j.jmr.2018.12.005.

[66] McCoy, K. M., Rogawski, R., Stovicek, O., McDermott, A. E., “Stability of nitroxide biradical TOTAPOL in biological samples,” Journal of Magnetic Resonance 303(2019), 10.1016/j.jmr.2019.04.013.

[67] Byeon, C.-H., Hansen, K. H., DePas, W., Akbey, Ü., “High-resolution 2D Solid-State NMR provides insights into Nontuberculous Mycobacteria,” Solid State Nuclear Magnetic Resonance 134(2024), 10.1016/j.ssnmr.2024.101970.

[68] Byeon, C.-H., Wang, Y.-H., Tunc, A., Franks, W. T., DePas, W. H., Akbey, Ü., “Structural Insights into Native Intact <em>Mycobacterium abscessus</em> by Conventional and Ultrahigh-field solid-state NMR at 1.2 GHz,” bioRxiv (2026), 10.64898/2026.05.19.726312.

[69] Ghosh, R., Xiao, Y., Kragelj, J., Frederick, K. K., “In-Cell Sensitivity-Enhanced NMR of Intact Viable Mammalian Cells,” J. Am. Chem. Soc. 143, no. 44 (2021), 10.1021/jacs.1c06680.

[70] Xiao, Y. L., Ghosh, R., Frederick, K. K., “In-Cell NMR of Intact Mammalian Cells Preserved with the Cryoprotectants DMSO and Glycerol Have Similar DNP Performance,” Frontiers in Molecular Biosciences 8(2022): 789478, 10.3389/fmolb.2021.789478.

[71] Flemming, H.-C., Wingender, J., Szewzyk, U., Steinberg, P., Rice, S. A., Kjelleberg, S., “Biofilms: an emergent form of bacterial life,” Nat Rev Microbiol 14, no. 9 (2016), 10.1038/nrmicro.2016.94.

[72] Hobley, L., Harkins, C., MacPhee, C. E., Stanley-Wall, N. R., “Giving structure to the biofilm matrix: an overview of individual strategies and emerging common themes,” FEMS Microbiology Reviews 39, no. 5 (2015), 10.1093/femsre/fuv015.

[73] Mishra, S., Huang, Y., Li, J., et al., “Biofilm-mediated bioremediation is a powerful tool for the removal of environmental pollutants,” Chemosphere 294(2022), 10.1016/j.chemosphere.2022.133609.

[74] Schooling, S. R., Beveridge, T. J., “Membrane Vesicles: an Overlooked Component of the Matrices of Biofilms,” Journal of Bacteriology 188, no. 16 (2006), 10.1128/jb.00257-06.

[75] Yordanov, N. D., Ranguelova, K., “Quantitative electron paramagnetic resonance and spectrophotometric determination of the free radical 4-hydroxy-2,2,6,6-tetramethylpiperidinyloxy,” Spectrochimica Acta Part A: Molecular and Biomolecular Spectroscopy 56, no. 2 (2000), 10.1016/S1386-1425(99)00248-6.

[76] Bobko, A. A., Kirilyuk, I. A., Grigor’ev, I. A., Zweier, J. L., Khramtsov, V. V., “Reversible reduction of nitroxides to hydroxylamines: Roles for ascorbate and glutathione,” Free Radical Biology and Medicine 42, no. 3 (2007), 10.1016/j.freeradbiomed.2006.11.007.

[77] Muratov, E., Keilholz, J., Kovács, Á. T., Moeller, R., “The biofilm matrix protects <i>Bacillu</i> <i>subtilis</i> against hydrogen peroxide,” Biofilm 9(2025), 10.1016/j.bioflm.2025.100274.

[78] Pierro, A., Bonucci, A., Normanno, D., et al., “Probing the Structural Dynamics of a Bacterial Chaperone in Its Native Environment by Nitroxide-Based EPR Spectroscopy,” Chemistry 28, no. 66 (2022), 10.1002/chem.202202249.

