## Supplementary material for "cMagnetic Resonance Reveals Radical Stability in Native Biofilms and its Role in Bacterial Resistance": Supplamentary Figures

**Table of Contents**

Experimental Details

Biofilm Sample Preparation S3

Cell Counting Assay S3

Biofilm Cell Viability Assay S3

EPR Sample Preparation S3

EPR Spectroscopy S4

ssNMR Spectroscopy S4

Electron Microscopy S4

Figure S1. Cell counting assay S6

Figure S2. Control experiments with 4-oxo-TEMPO S7

Figure S3. AsymPolPOK reduction over 24 hours S8

Figure S4. Reduced AsymPolPOK and monoradical CW-EPR spectra S9

Figure S5. AsymPolPOK biradical/monoradical simulations S10

Figure S6. AsymPolPOK DNP signal reduction S11

Figure S7. WT biofilm, 4-oxo-TEMPO, and oxidizing agents S12

Figure S8. CW-EPR intensity versus doubly integrated intensity S14

Figure S9. CW-EPR linewidth analysis S15

Figure S10. Non-normalized CW-EPR time scans S16

Figure S11. 80 K CW-EPR time scans S17

Figure S12. CW-EPR spectra of biofilm without radical S17

References S18

**Biofilm Sample Preparation.** *Pseudomonas fluorescens* Pf0-1 colony biofilms were prepared as previously described.[1-4] Briefly, liquid cultures were spotted onto solid Pseudomonas Agar F (PAF, Difco) plates and incubated at 30 °C for 72 h. Biofilms were gently scraped from the agar surface and stored at 4 °C in microcentrifuge tubes. Planktonic samples were prepared under the same conditions except that the cells were grown overnight in liquid PAF without agar, shaking at 200 RPM, and centrifuged for 5 minutes at 3,000 x *g*. For $^{13}C$-detected ssNMR, samples were packed into thin-walled 3.2 mm MAS rotors with 40–60 mg of material. Wet biofilms were filled into rotors using pipettes and benchtop centrifugation. Due to the high water content of native samples, the dry biofilm mass is low. Drying at ~50 °C under ambient conditions increased the solid content by up to tenfold. Both hydrated and dried biofilm samples were analyzed.

**Cell Counting Assay.** Wild-type Pf0-1 was grown overnight in PAF broth with shaking (200 rpm, 30°C) for planktonic cultures or spotted (20 µl) onto PAF plates and incubated statically at room temperature for 72 hours for biofilm cultures. For CFU enumeration, planktonic cells were harvested by centrifugation of 1 ml culture and resuspension in 1 ml PBS; a parallel 30 µl aliquot was taken directly without centrifugation as a viability control. Biofilm cells were scraped individually (one spot per biological replicate) into 1 ml PBS. All samples were serially diluted 10-fold in PBS and plated in duplicate. CFU/ml was calculated from the best countable dilution. Cell density (cells/mg) was calculated by multiplying CFU/ml by the total sample volume and dividing by pellet weight. Biofilm pellets had approximately 2-fold higher cell density than planktonic ($p = 0.003$, unpaired t-test), and cell numbers were normalized for EPR experiments by taking 10 mg of planktonic and 5 mg of biofilm cells, each delivering $\sim 4.3 \times 10^8$ cells. Centrifugation did not significantly affect cell viability (mean ~100.5%, one-sample t-test vs 1, $p = 0.89$). Data are presented as mean ± SEM, statistical analyses were performed in GraphPad Prism.

**Biofilm Cell Viability Assay**. WT Pf0-1 colony biofilms were prepared by spotting overnight cultures onto PAF plates in 20 µL drops and incubated at room temperature for 3 days. One colony was scraped into a tube via inoculation loop for each condition and timepoint (done in biological triplicate for each). Three conditions at two timepoints (10 min & 60 min exposure): 1) Pellet incubated in refrigerator (4 °C) and 2) Pellet exposed to liquid nitrogen. After exposure, conditions 1 and 2 were resuspended in 1 mL PBS. All samples were diluted to $10^{-7}$, and 50 µL of each dilution (including undiluted) were plated in triplicate onto PAF plates and incubated at 30 °C overnight, then isolated colonies were counted. The effect of refrigerator and liquid-nitrogen treatment on the cell-viability of the biofilm was determined by a two-way ANOVA and Tukey's test.

**EPR Sample Preparation**. Frozen biofilm stocks were thawed, vortexed, and briefly centrifuged at a low speed to bring all material to the bottom of the tube. For the samples in Figures 2 and 4, approximately 10 mg of biofilm sample was taken into a clean microcentrifuge tube. In separate tubes, 50 µL solutions of 10 mM 4-oxo-TEMPO (MW: 170 g/mol) and 10 mM AsymPolPOK (MW: 765 g/mol) in 10% (v/v) DMSO in water were prepared. For the samples in Figures 2 and 3, the volume-to-cell mass ratio is provided in the main text. In the case of hydrogen peroxide or potassium ferricyanide, the 50 μL solutions of varying concentrations were prepared before adding to the 10 mg of biofilm sample. Once added to the biofilm, solutions were quickly vortexed to make the biofilm sample more homogeneous and then drawn into Pyrex capillary tubes (1 mm outer diameter). Capillary tubes were sealed and placed into EPR sample tubes (3 mm outer diameter). EPR measurements were performed within 5 minutes of mixing with the radical solution. The amounts are approximate due to varying amounts of biofilm mass in the samples due to different hydration levels and cell counts.

**EPR Spectroscopy**. Continuous-wave (CW) EPR experiments were performed on the Bruker ElexSys E680 FT/CW spectrometer with a Bruker ER4122 SHQE-W1 high-resolution resonator at X-band (~9.8 GHz) frequencies. Spectra were acquired at room temperature and 80 K at a center field of 3512 G with a sweep width of 150 G. A total of 1024 data points were collected using a conversion time of 20.48 ms. A modulation frequency of 100 kHz and a modulation amplitude of 1 G were used. For time-resolved EPR experiments, spectra were collected within a range of every 30 seconds to 10 minutes, depending on the sample.

**ssNMR Spectroscopy**. All ssNMR experiments were recorded on a Bruker Avance III 750 MHz (17.6 T) spectrometer using a low-temperature triple-resonance (HCN) 3.2 mm MAS probe. Experiments were performed at a magic-angle spinning (MAS) rate of 10 kHz and 275 K set temperature, corresponding to ambient sample temperature. $^{1}$H and $^{13}$C pulses were 3.3 ms and 5 ms, respectively. Cross-polarization (CP) experiments employed a 1 ms contact time and a 70–100% ramp on the $^{1}$H channel. SPINAL-64 decoupling was applied at ~90 kHz during acquisition. $^{1}$H chemical shifts were externally referenced to DSS (0 ppm), and the $^{13}$C chemical shifts were indirectly referenced.[5] One-dimensional $^{13}$C INEPT and CP spectra were acquired with 36 k scans, using recycle delays of 1 s, with total acquisition times of approximately ~10 h. One-dimensional spectra were apodized using a Gaussian window function (GB = 0.05, LB = 35 Hz). Two-dimensional (2D) $^{1}$H-detected $^{1}$H–$^{13}$C INEPT spectra for wet biofilms with/without TEMPO radical were acquired with 196 transients. Two different TEMPO concentrations were used to see the concentration dependent radical effect, with 10 mM and excess TEMPO in the biofilm samples. 1 s recycle delay and 128 $t_1$ increments ($\Delta t_1$ = 38 ms) were used, resulting in total experimental times of ~7 h. The 2D spectrum was processed using Gaussian apodization in the direct dimension (GB = 0.1, LB = -35 Hz) and a sine-squared window function (SSB = 3) in the indirect dimension. The details of the 2D $^{13}$C-detected $^{1}$H–$^{13}$C INEPT spectra for wet biofilm with tentative assignments were given in our previous work. [6] The DNP ssNMR spectra were acquired as previously described.[7] A 3.2 mm LT DNP probe is utilized at ~100 K and 10 kHz MAS using a sapphire rotor with ~30 mg of sample on a 600 MHz Bruker NEO ssNMR spectrometer. The 1D $^{13}$C CP ssNMR spectra were recorded with 256 scans with ~3.5 seconds recycle delay with/without microwave irradiation to estimate DNP signal enhancement.

**Electron Microscopy**. Negative-staining electron microscopy (EM) micrographs of wild-type (WT) planktonic cells, WT biofilm, mucoid biofilm, and dry biofilm. Strains were recorded on a Tecnai TF20 microscope (Thermo Fisher Scientific, MA) with a field emission gun operating at

200 kV equipped with a TVIPS XF416 complementary metal-oxide semiconductor camera (TVIPS GmbH, Gilching, Germany). Four hundred mesh copper grids with a continuous carbon film were glow discharged for 60 s. 3 μL of sample was applied to grids and incubated for 10 s and side blotting on filter paper. Staining was done with 2% (w/v) uranyl acetate for 10 s and side blotted on filter paper.

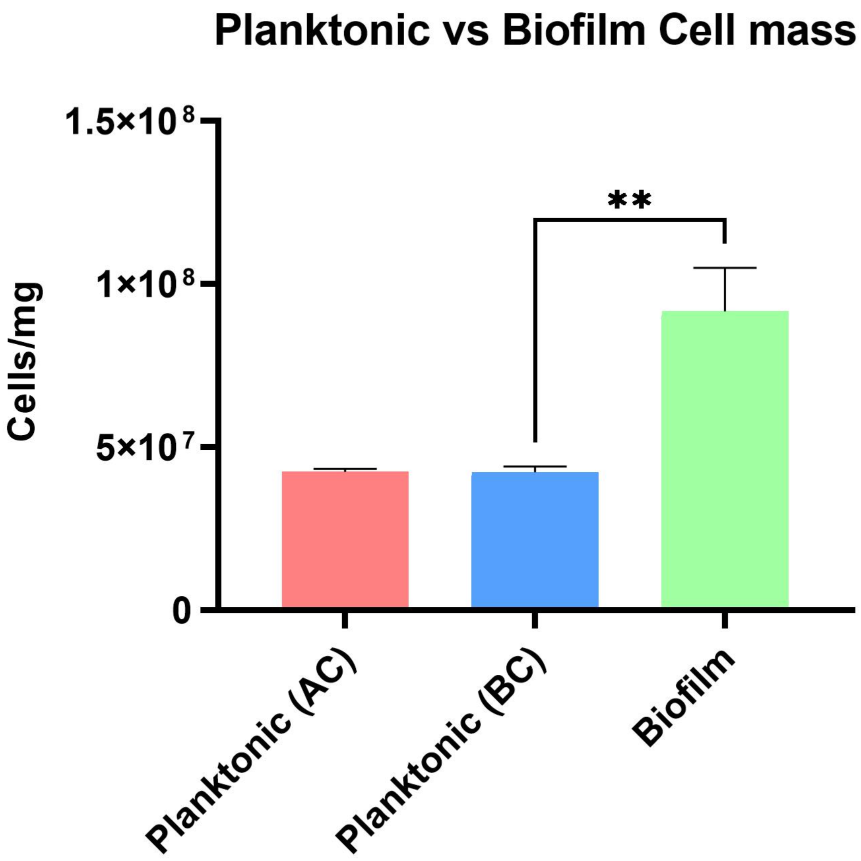


**Figure S1**. **Biofilm cells exhibit higher cell density per mg than planktonic cultures.** Wild-type Pf0-1 was grown either planktonically in broth overnight with shaking (200 rpm, 30°C) (before centrifugation, BC; after centrifugation, AC) or as surface-associated biofilms on solid agar for 72 hours at room temperature. Cell density was quantified by CFU enumeration and normalized to pellet weight (cells/mg). Biofilm cultures showed approximately 2-fold higher cell density compared to planktonic cultures. Accordingly, 5 mg of biofilm and 10 mg of planktonic cells were used to deliver equivalent cell numbers (~$4.3 \times 10^8$ cells) for EPR experiments. Bars represent mean ± SEM (n = 3). Statistical significance was determined by unpaired two-sample t-test (* $p < 0.05$).

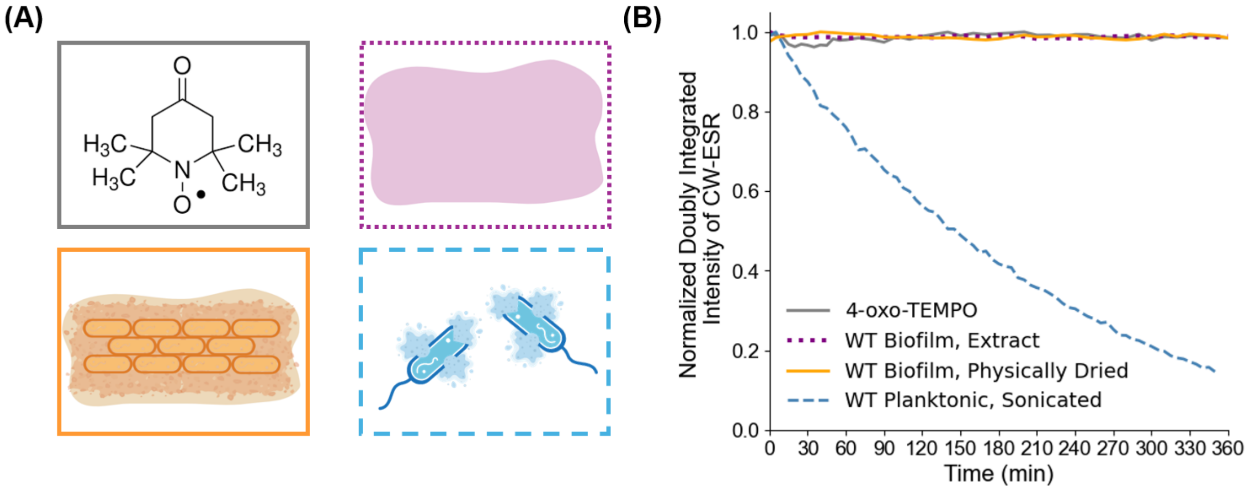


**Figure S2.** (A) Illustrations of the samples shown in (B). (B) Doubly integrated intensity of continuous-wave EPR spectra as a function of time. Control experiments of 4-oxo-TEMPO radical (solid grey), ECM extract from WT biofilm (dotted purple), physically dried WT biofilm (solid orange), and sonicated WT planktonic cells (dashed light blue).

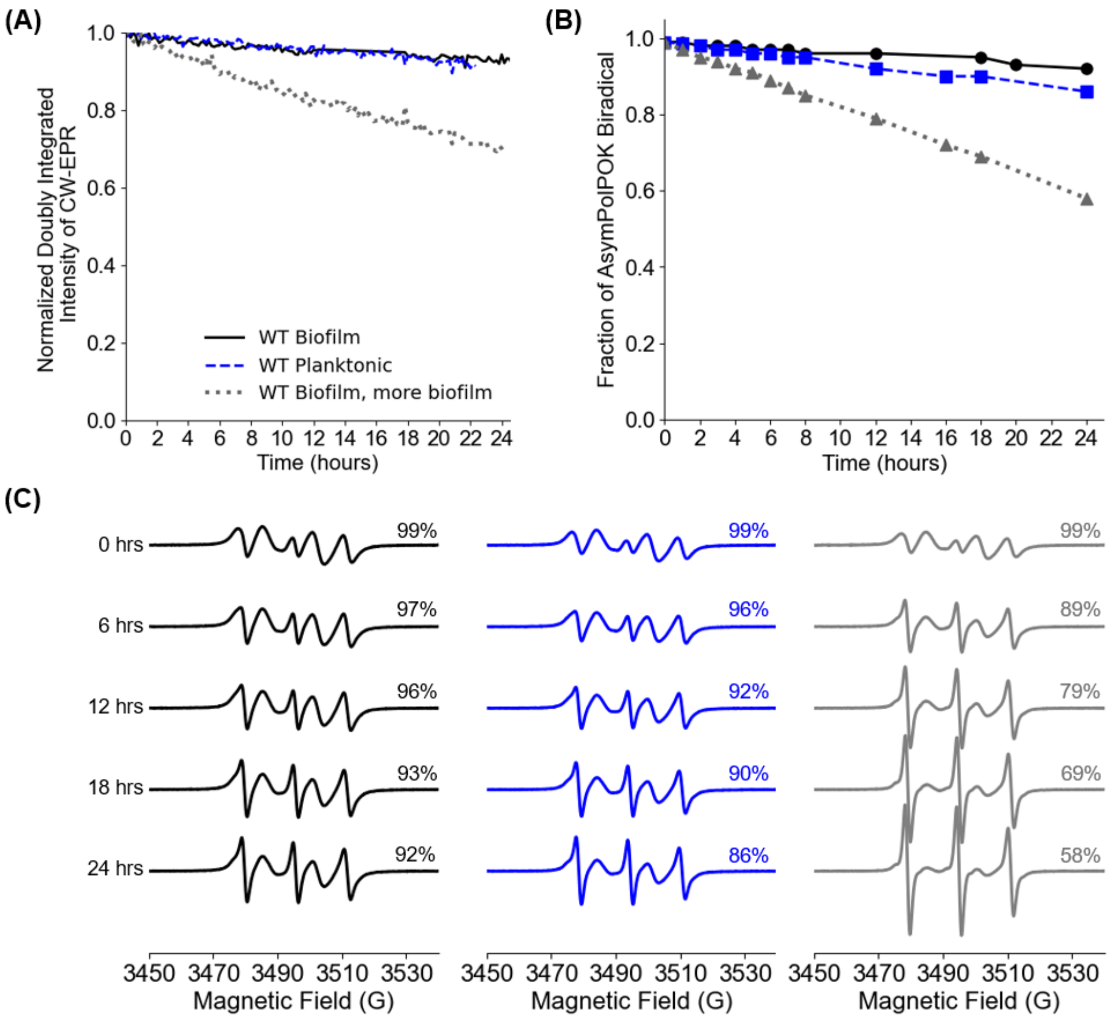


**Figure S3**. 24-hour data for AsymPolPOK with WT biofilm, planktonic cells, and higher concentration of WT biofilm. Further explanation of the simulation method is in Figure S5.

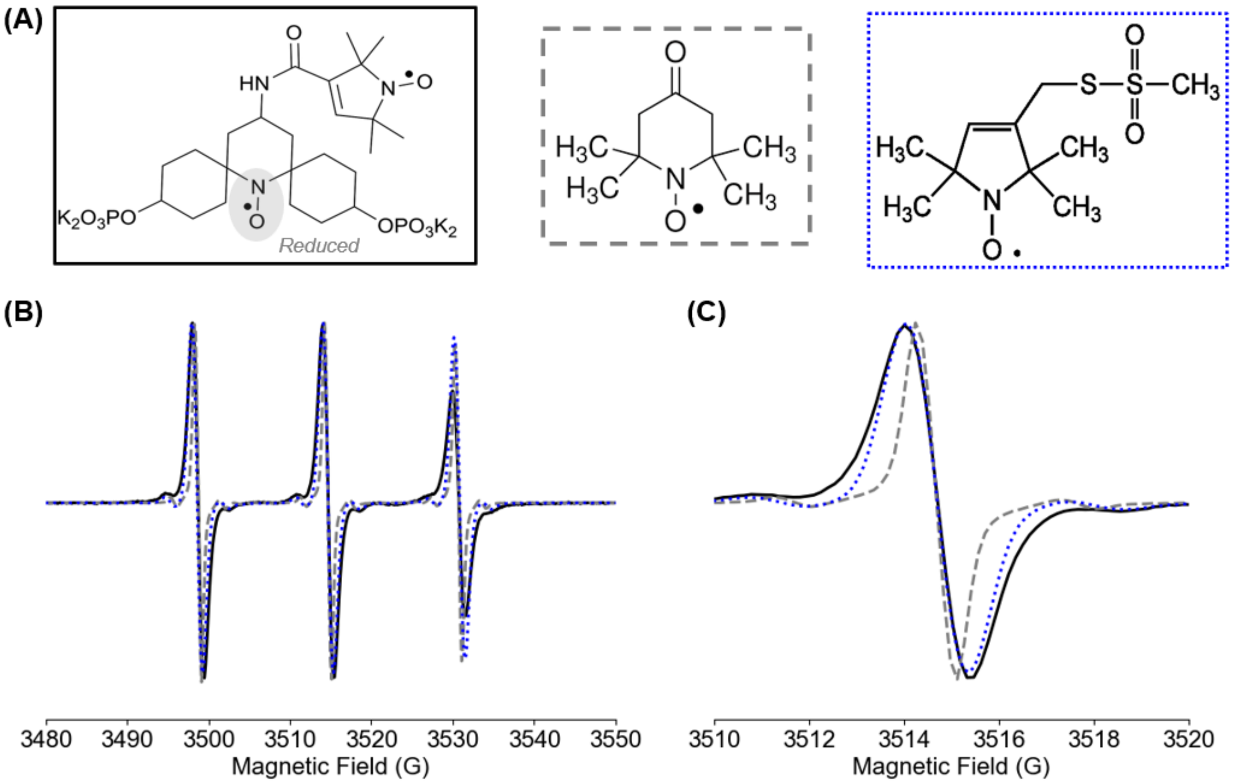


**Figure S4**. Comparison of room temperature CW-EPR spectra for nitroxide-based radicals and reduced AsymPolPOK. (A) Structures of the radicals studied. The biradical AsymPolPOK was reduced to a monoradical. The single nitroxide radicals used were 4-oxo-TEMPO (dashed grey) and (1-oxyl-2,2,5,5-tetramethylpyrroline-3-methyl)methanethiosulfonate spin label (MTSSL) (dotted blue). (B) CW-EPR spectra. The AsymPolPOK spectrum (solid black) was collected on 10 mM AsymPolPOK solution after 48 hours of incubation with WT biofilm. The 4-oxo-TEMPO and MTSSL spectra were acquired on fresh 10 mM solutions of each. The hyperfine splitting $A_{iso}$ was nearly same for each sample (AsymPolPOK: 15.9 G, 4-oxo-TEMPO: 16.0 G, MTSSL: 16.1 G) (C) Middle peak of the CW-EPR spectra to highlight differences in peak-to-peak linewidths (AsymPolPOK: 1.3 G, 4-oxo-TEMPO: 0.9 G, MTSSL: 1.3 G).

After reduction of AsymPolPOK to a monoradical, the spectral linewidth most closely matches that of the 5-membered MTSSL nitroxide. Based on this, we suggest that the 6-membered radical is reduced preferentially.

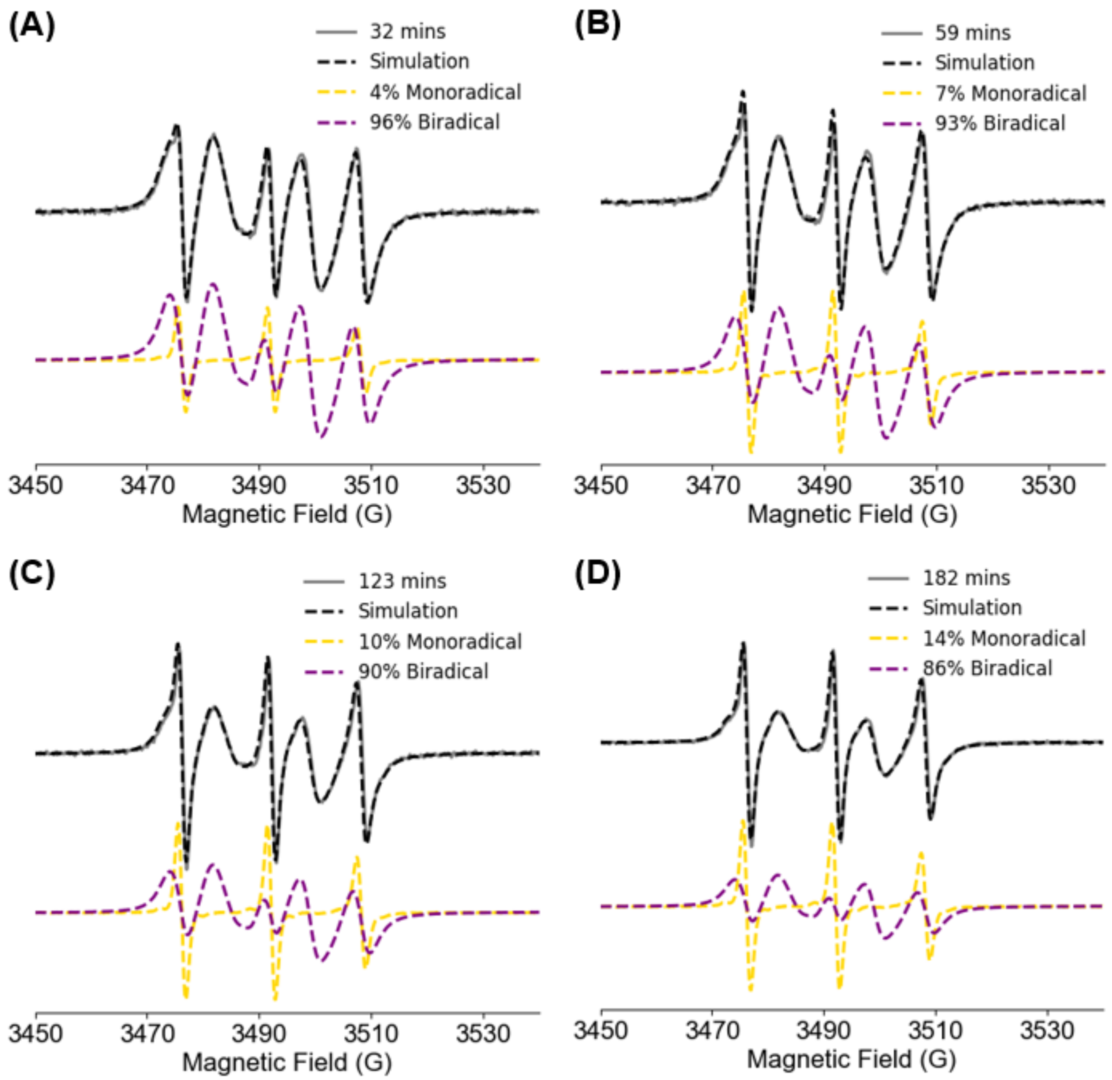


**Figure S5**. Representative EPR spectra simulations of AsymPolPOK reduction over time. Each experimental spectrum (solid grey) was collected at different points during the reduction of the radical. Simulations (dashed black) were performed by normalizing the approximate double integral of the biradical (dashed purple) and monoracial (dashed gold) spectra, weighting them accordingly, then adding them together until there was a match to the experimental spectra. The biradical spectrum was collected as free AsymPolPOK in solution, and the monoradical spectrum was collected after incubating AsymPolPOK with biofilm for 48 hours.

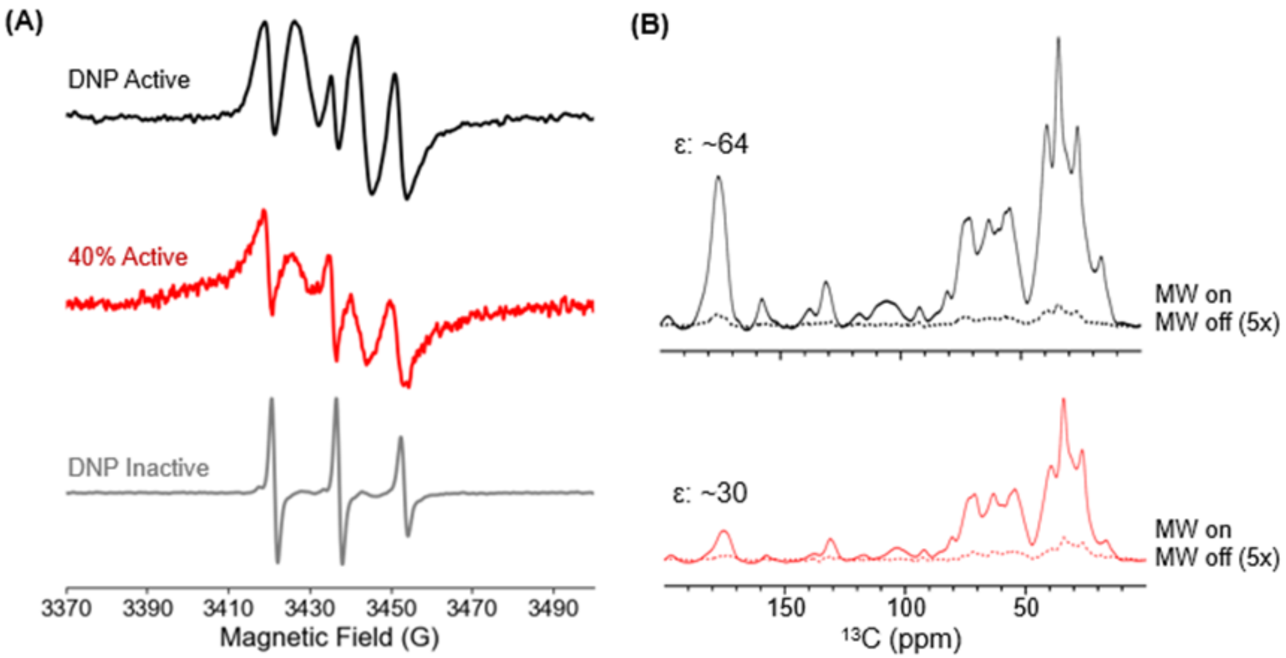


**Figure S6.** (A) EPR spectra of AsymPolPOK in biofilm from initial incubation ("DNP Active", black) to partial reduction ("40% Active", red) and final form ("DNP Inactive", grey). Each spectrum is normalized to its maximum intensity. (B) $^{13}C$ DNP-enhanced ssNMR spectra of Active and 40% Active samples. The microwave on and off spectra are shown as solid and dashed lines, respectively.

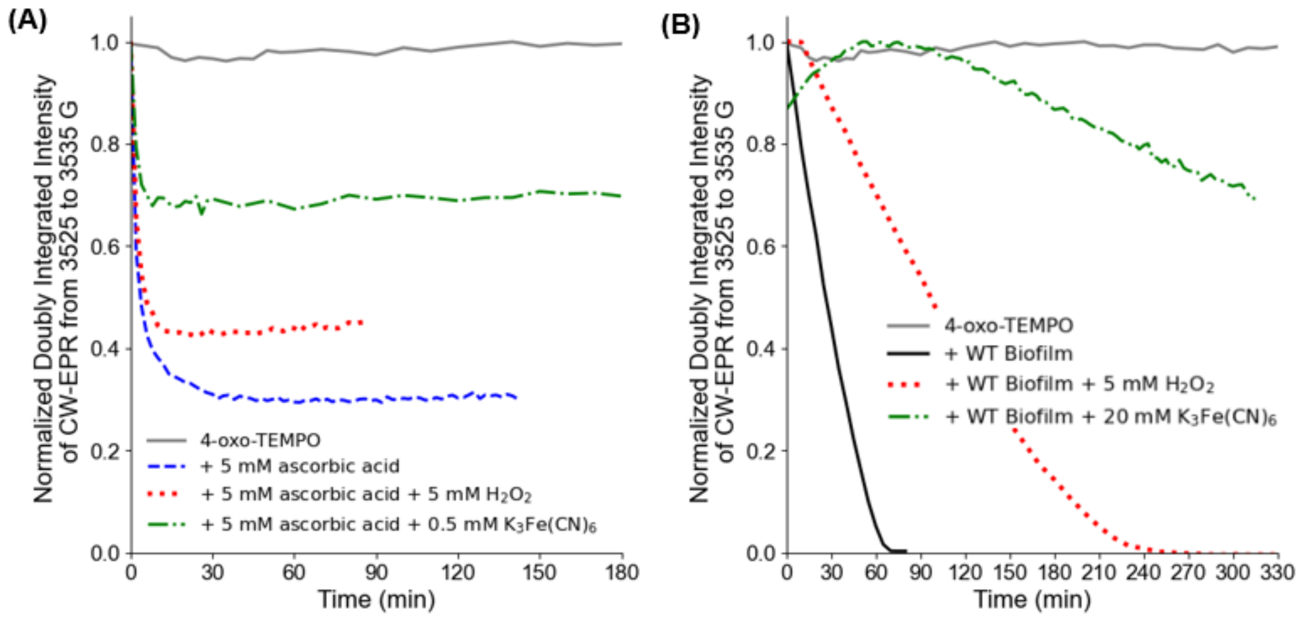


**Figure S7.** (A) Doubly integrated intensity of continuous-wave EPR spectra as a function of time for 10 mM 4-oxo-TEMPO (solid grey), 10 mM TEMPO with 5 mM ascorbic acid (dashed blue), 10 mM TEMPO with 5 mM ascorbic acid and 5 mM hydrogen peroxide (dotted red), and 10 mM TEMPO with 5 mM ascorbic acid and 0.5 mM $K_3Fe(CN)_6$. The x-axis is from 0 to 180 minutes to highlight the sharp decrease in the signal upon adding a reducing agent. (B) CW-EPR signal as a function of time for radical (solid grey), WT biofilm (solid black), WT biofilm with TEMPO and 5 mM hydrogen peroxide (dotted red), and WT biofilm with TEMPO and 20 mM $K_3Fe(CN)_6$ (dash dotted green).

Given the inherent reductive environments shown in biofilm samples, we next investigated strategies to prolong radical lifetimes in those samples, which is an essential requirement for effective DNP applications. One approach to prevent radical inactivation is rapid sample preparation followed by immediate freezing, although this method compromises molecular mobility and results in broader ssNMR resonances. Nevertheless, freezing of biological samples in DNP ssNMR is routinely applied and offers a more quantitative composition analysis since both the rigid and the flexible signals are detected at once due to freezing. Alternatively, the addition of an oxidizing agent offers a promising means of stabilizing radicals in situ.[8-9]

To explore this, we examined the reduction kinetics of 4-oxo-TEMPO by ascorbic acid, a common intracellular reductant well-studied by EPR.[9-10] **Figure S7A** shows that ascorbic acid induces a rapid decay of the EPR signal within minutes, followed by equilibration after approximately 20 minutes at which point all of the ascorbate is consumed. When the oxidizing agent, hydrogen peroxide ($H_2O_2$), is added to ascorbic acid at equimolar concentrations, the radical is much less reduced, and the overall EPR signal remains higher than in the absence of peroxide. These findings suggest that oxidant supplementation can partially counteract the reductive quenching of nitroxides, offering a viable strategy to preserve radical activity within biofilms.

However, excessive concentrations of hydrogen peroxide are known to be cytotoxic,[11] making it nonideal for biophysical characterization of native samples. Alternatively, potassium ferricyanide ($K_3Fe(CN)_6$) has been shown to increase the nitroxide lifetime in cells and cell extracts and offers a more biocompatible solution.[8, 12-14] **Figure S7A** shows that with 0.5 mM potassium ferricyanide, the EPR signal is static at a higher intensity than with a ten-fold higher hydrogen peroxide concentration. These findings altogether show that potassium ferrocyanide is a very potent chemical to prevent radical deactivation in native biofilms compared to $H_2O_2$ and can result in high efficiency DNP ssNMR applications.

We further investigated the impact of the oxidants on the reduction kinetics of TEMPO within the WT biofilm. Remarkably, the addition of 5 mM hydrogen peroxide significantly slows the decay of the nitroxide radical, as shown in **Figure S7B**. Specifically, the presence of hydrogen peroxide extended the lifetime of the EPR signal by almost four-fold compared to the untreated biofilm. **Figure S7B** shows that 20 mM potassium ferricyanide increases the lifetime of the nitroxide signal, and we do not observe reduction beyond 20% within 5-6 hours of monitoring. We attribute the increase in the signal at the early timepoints to the reoxidation of any initially reduced TEMPO. Alternatively, a locally high concentration of radicals, due to localization in specific biofilm regions, would initially cause line broadening and reduced signal intensity. As the experiment progresses, the radical concentration would decrease, diminishing hyperfine coupling and leading to narrower linewidths and increased signal intensity up to approximately 60-90 minutes (**Figure S8-10**). Beyond this point, the signal would begin to decline again as the overall radical population continues to decrease. Finally, the rapid freezing of the biofilm sample, as an alternative method to slow the reduction of TEMPO, does not result in a reduction of the EPR signal over 6 hours (**Figure S11**). This indicates that at liquid nitrogen temperatures, diffusion is absent and interactions with the radical cannot occur. Overall, these results indicate that oxidizing agents, in particular potassium ferricyanide, can effectively inhibit radical quenching in native biofilm environments and thus potentially preserve DNP enhancement during ssNMR measurements. However, additional experiments are needed to assess the impact of oxidizing agents on biofilm viability.

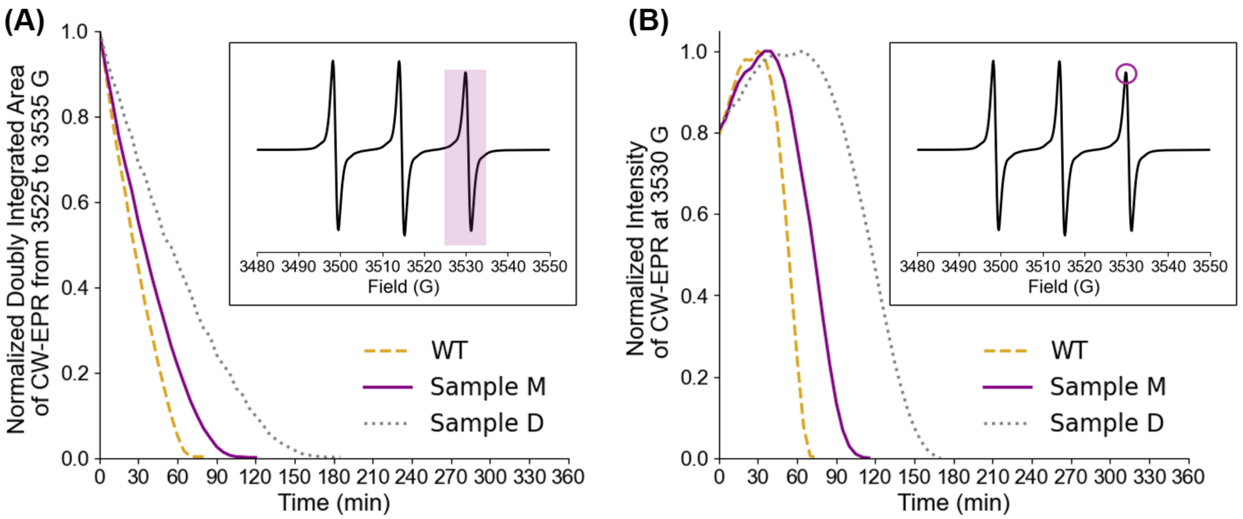


**Figure S8**. (A) Doubly integrated intensity of room temperature CW-EPR spectra as a function of time for 4-oxo-TEMPO in WT biofilm (dashed gold), mucoid biofilm (solid purple, M) and dry biofilm (dotted grey). The inset shows the spectrum, and the highlighted region is the bounds of integration (3525 – 3535 G). (B) Normalized intensity of room temperature CW-EPR spectrum as a function of time at 3530 G. The inset shows the peak where the intensity was recorded.

The increase in the intensity in the beginning of the time course is a surprising finding, as one would expect an overall decrease in nitroxide EPR signal due to the cellular environment. The increase in intensity is due to a narrowing of the spectral lineshape over time, and this behavior as a function of time has been observed in in-cell EPR experiments by Pierro et al.[15] They attributed the sharpening to an increase in label mobility which could be explained by possible protease action of protein unfolding. In their case, a spin-labeled protein could be fragmented by a protease, causing a region with the spin label to become more mobile. In our work, it is possible that the nitroxide interacts in a region of the biofilm that becomes more flexible with the interaction, leading to an increase in mobility. It is difficult to exactly determine the cause of the narrowing, but a protein or peptide unfolding is a possible hypothesis.

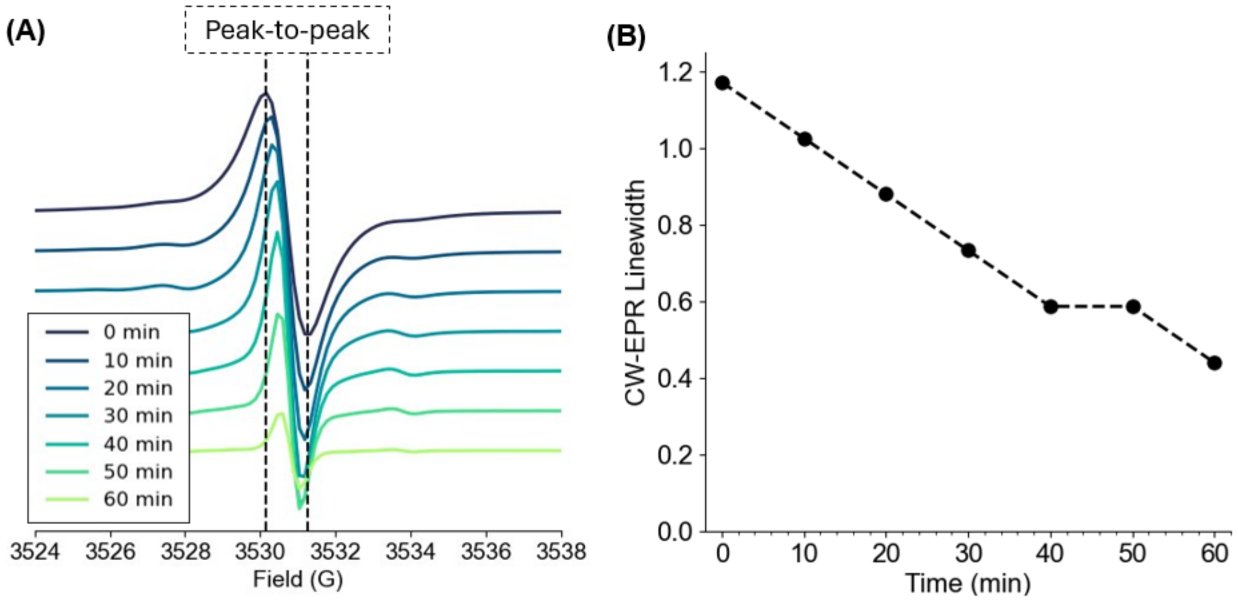


**Figure S9**. (A) CW-EPR spectra of the WT biofilm sample with 4-oxo-TEMPO from 3524 to 3538 G to highlight the third hyperfine peak. The color gradient shows the change in the peak-to-peak linewidth in the EPR spectrum as a function of time. (B) Plot to show the CW-EPR linewidth as a function of time.

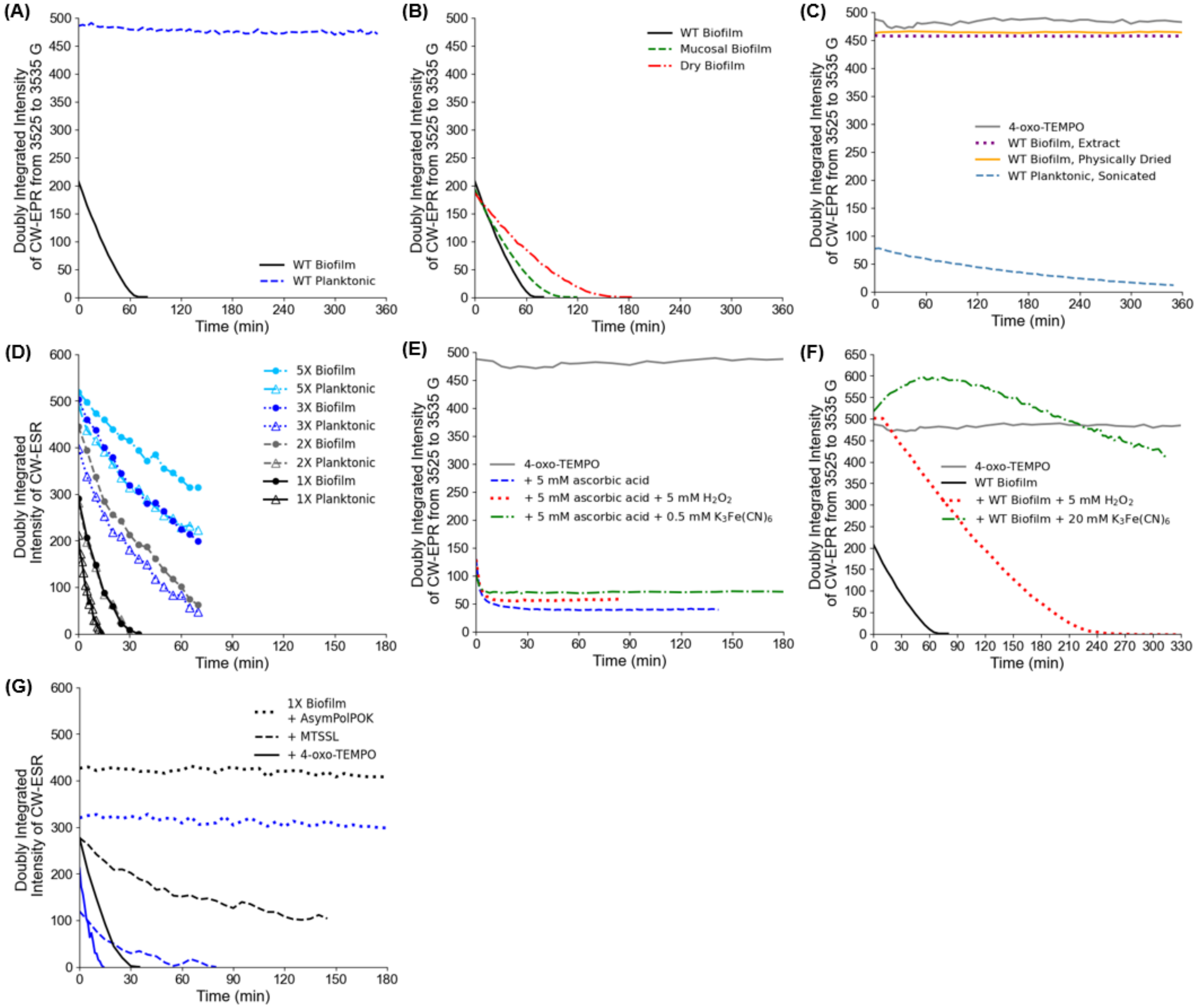


**Figure S10**. Non-normalized CW-EPR signal decays from the Main Text and SI.

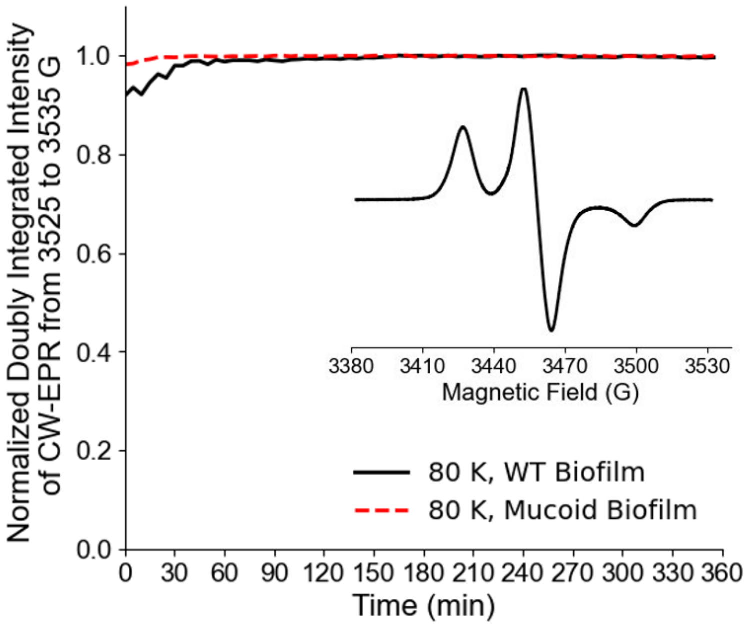


**Figure S11**. Doubly integrated intensity of 80 K continuous-wave EPR spectra as a function of time for WT biofilm (black solid) and mucoid biofilm (red dashed). The inset is the 80 K CW-EPR spectrum for the WT biofilm at time 0.

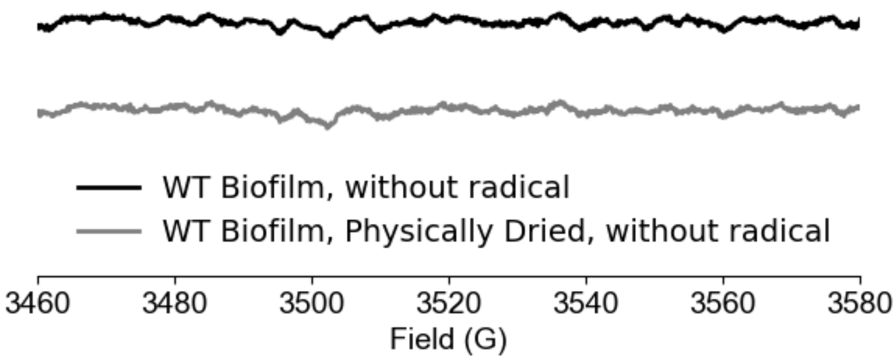


**Figure S12**. CW-EPR spectra of 10 mg of WT biofilm in 10% DMSO (black) and physically dried WT biofilm (grey) without the addition of 4-oxo-TEMPO. The grey spectrum is staggered below the black spectrum for visualization.

**References**


[1] A. F. Evans, M. K. Wells, J. Denk, W. Mazza, R. Santos, A. Delprince, W. Kim, *Journal of Bacteriology* **2022**, *204*, e00285–00222.

[2] W. Kim, S. B. Levy, K. R. Foster, *Nat Commun* **2016**, *7*, 10508.

[3] W. Kim, F. Racimo, J. Schluter, S. B. Levy, K. R. Foster, *Proceedings of the National Academy of Sciences* **2014**, *111*, E1639–E1647.

[4] W. P. J. Smith, Y. Davit, J. M. Osborne, W. Kim, K. R. Foster, J. M. Pitt-Francis, *Proceedings of the National Academy of Sciences* **2017**, *114*, E280–E286.

[5] D. S. Wishart, C. G. Bigam, A. Holm, R. S. Hodges, B. D. Sykes, *J Biomol NMR* **1995**, *5*, 67–81.

[6] C.-H. Byeon, T. Kinney, H. Saricayir, S. Srinivasa, M. K. Wells, W. Kim, Ü. Akbey, *Journal of Magnetic Resonance* **2023**, *357*, 107587.

[7] C.-H. Byeon, T. Kinney, H. Saricayir, K. H. Hansen, F. Scott, S. Srinivasa, M. K. Welss, F. Mentink-Vigier, W. Kim, U. Akbey, *Angewandte Chemie* **2025**, e202418146.

[8] K. Singewald, M. J. Lawless, S. Saxena, *Journal of Magnetic Resonance* **2019**, *299*, 21–27.

[9] N. D. Yordanov, K. Ranguelova, *Spectrochimica Acta Part A: Molecular and Biomolecular Spectroscopy* **2000**, *56*, 373–378.

[10] A. A. Bobko, O. V. V. Efimova, Maxim A., V. V. Khramtsov, *Free Radical Research* **2012**, *46*, 1115–1122.

[11] E. Muratov, J. Keilholz, Á. T. Kovács, R. Moeller, *Biofilm* **2025**, *9*, 100274.

[12] M. J. Lawless, A. Shimshi, T. F. Cunningham, M. N. Kinde, P. Tang, S. Saxena, *ChemPhysChem* **2017**, *18*, 1653–1660.

[13] K. M. McCoy, R. Rogawski, O. Stovicek, A. E. McDermott, *Journal of Magnetic Resonance* **2019**, *303*, 115–120.

[14] P. Wipf, J. Xiao, J. Jiang, N. A. Belikova, V. A. Tyurin, M. P. Fink, V. E. Kagan, *J. Am. Chem. Soc.* **2005**, *127*, 12460–12461.

[15] A. Pierro, A. Bonucci, D. Normanno, M. Ansaldi, E. Pilet, O. Ouari, B. Guigliarelli, E. Etienne, G. Gerbaud, A. Magalon, V. Belle, E. Mileo, *Chemistry* **2022**, *28*, e202202249.
